# Principles that govern competition or co-existence in Rho-GTPase driven polarization

**DOI:** 10.1101/148064

**Authors:** Jian-geng Chiou, Timothy C. Elston, Thomas P. Witelski, David G. Schaeffer, Daniel J. Lew

## Abstract

Rho-GTPases are master regulators of polarity establishment and cell morphology. Positive feedback enables concentration of Rho-GTPases into clusters at the cell cortex, from where they regulate the cytoskeleton. Different cell types reproducibly generate either one (e.g. the front of a migrating cell) or several clusters (e.g. the multiple dendrites of a neuron), but the mechanistic basis for uni-polar or multi-polar outcomes is unclear. The design principles of Rho-GTPase circuits are captured by reaction-diffusion models based on conserved aspects of Rho-GTPase biochemistry. Some such models display rapid winner-takes-all competition between clusters, yielding a unipolar outcome. Other models allow prolonged co-existence of clusters. We derive a “saturation rule” general to all relevant models that governs the timescale of competition, and thereby predicts whether the system will generate uni-polar or multi-polar outcomes. We suggest that the saturation rule is a fundamental property of the Rho-GTPase polarity machinery, regardless of the specific feedback mechanism.

## Introduction

Complex cell morphologies arise, in part, through the specialization of cortical domains (e.g., the apical and basal domains of epithelial cells, or the front and back of migratory cells). Elaboration of such domains involves the local accumulation of active Rho-family GTPases, which regulate cytoskeletal elements to promote specific downstream events, such as vesicle trafficking, membrane deformation, or directed growth (***Caceres et al., 2012***; ***Etienne-Manneville and Hall, 2002***; ***Yang, 2008***). For some cells, it is vital to establish a single specialized domain (e.g. the front of a migrating cell), whereas others require the establishment of multiple domains simultaneously (e.g. the dendrites of a neuron) (***Dotti et al., 1988***; ***Wu and Lew, 2013***). The mechanistic basis for specifying uni- or multi-polar outcomes remains elusive.

Rho-family GTPases switch between GTP-bound active and GDP-bound inactive forms (*Figure 1*A). Active GTPases are tethered to the inner surface of the plasma membrane, where diffusion is slow. In contrast, inactive GTPases are preferentially bound by guanine nucleotide dissociation inhibitors (GDIs), which extract the bound GTPase to the cytoplasm, where their diffusion is comparatively fast. Activated GTPases can promote local activation of cytosolic GTPases via positive feedback. This generates a membrane domain with concentrated active GTPase, concomitantly depleting the cytosolic GTPase pool (*Figure 1*B). Synthesis and degradation of GTPases occurs on a slow timescale compared to activation and inactivation (for example, in budding yeast the Rho-GTPase of Cdc42 polarizes within 2 minutes but has a half-life of more than 20 hours) (***Gladfelter et al., 2001***; ***Howell et al., 2009***; ***Wedlich-Soldner et al., 2004***). Thus, the general dynamics of the system can be captured by mass-conserved activator-substrate (MCAS) models, with a slowly-diffusing activator and a rapidly-diffusing substrate (*Figure 1*C) (***Goryachev and Pokhilko, 2008***; ***Mori et al., 2008***; ***Otsuji et al., 2007***). Such models can generate local peaks of activator, reflecting the establishment of a polarized concentration profile of active GTPase (*Figure 1*D).

**Figure 1.**
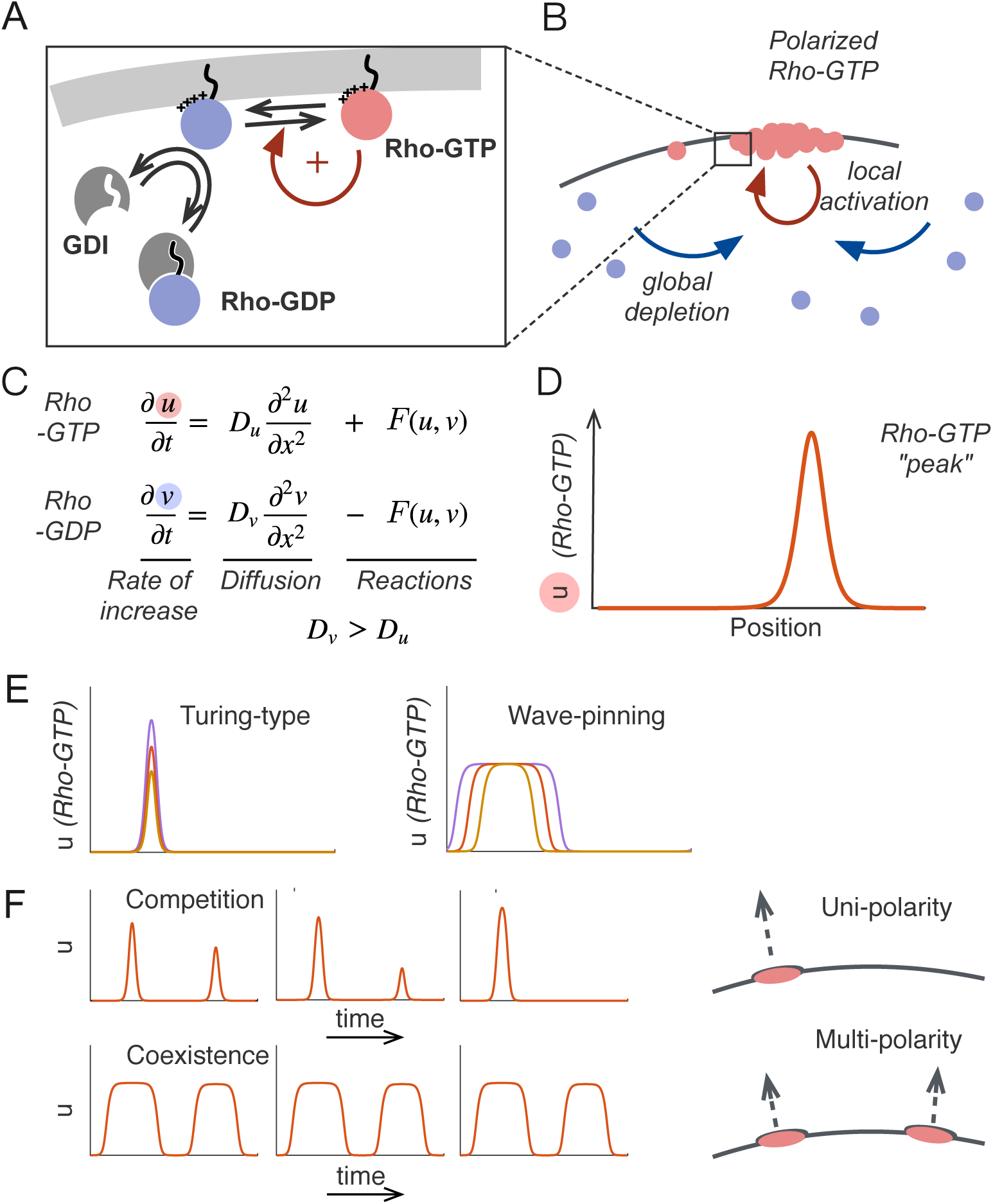
Polarity establishment and competition in mass conserved activator-substrate (MCAS) models. A) Rho-GTPases are tethered to the plasma membrane by prenylation. The inactive GDP-bound form, or “substrate”, is preferentially bound by the GDI, masking the prenyl group and extracting the substrate to the cytoplasm. The active GTP-bound form, or “activator”, promotes local activation of more substrate, yielding positive feedback. B) Local activation via positive feedback and depletion of the substrate in the cytosol generates an activator-enriched domain on the cortex. C) The interconversions of Rho-GTPases between active and inactive forms can be modeled as a system of two reaction-diffusion equations governing the dynamics of the slowly-diffusing activator *u* and the rapidly-diffusing substrate *v*. The model is mass-conserved: generation of *u* is precisely matched by consumption of *v* (and vice versa) in the reaction term *F*(*u, v*). D) MCAS models generate peaks in the profile of *u*, representing concentrated active Rho-GTPase on the membrane. E) Turing-type models (***Equation 4***) can generate sharp peaks of different heights, while wave-pinning models (***Equation 5***) can generate flat-topped mesas of different width, when total Rho-GTPase content *M* increases. *M* = 4, 6, 10 for Turing-type model and *M* = 30, 40, 50 for wave-pinning model. F) When two peaks of unequal size form in Turing-type models, they compete rapidly and resolve to a single peak, whereas two mesas of unequal size in Wave-pinning models are meta-stable. Parameter values are *a* = 1 *μm*^2^, *b* = 1 *s*^−1^ and *D_u_* = 0.01 *μm*^2^ *s*^−1^, *D_v_* = 1 *μm*^2^ *s*^−1^ for both models, and *k* = 1 *μm*^2^ for wave-pinning model. All models were simulated on domain size *L* = 10 *μm*.

Proposed MCAS models differ primarily in the formulation of the positive feedback mechanism. One set of models yields Turing instability (***Goryachev and Pokhilko, 2008***; ***Otsuji et al., 2007***), where positive feedback is sufficient to amplify molecular-level fluctuations leading to peak formation. Classically, Turing systems can generate single or multiple peaks (***Gierer and Meinhardt, 1972***; ***Turing, 1952***), depending on whether the size of the modeled domain is larger than a characteristic wavelength dependent on the reaction and diffusion parameters. However, even when multiple peaks emerge from the homogeneous state, most of the peaks in Turing-type MCAS models eventually disappear through a process called “competition”, leaving a single large peak as the winner (***Howell et al., 2012***; ***Otsuji et al., 2007***; ***Wu et al., 2015***). ***Otsuji et al.*** (***2007***) reasoned that competition arose due to mass-conservation, and further suggested that this might be a general behavior of Turing-type MCAS models. In biological systems, competition-like behavior was observed during polarity establishment in yeast cells, where it was suggested to underlie the growth of only one bud per cell cycle (***Howell et al., 2012***, ***2009***; ***Wu et al., 2015***).

Another set of models relies on bistable reaction kinetics to produce “wave-pinning” behavior (***Beta et al., 2008***; ***Mori et al., 2008***, ***2011***; ***Ozbudak et al., 2005***). Such models can generate membrane domains with separate phases of uniform high or low activator concentrations connected by a sharp “wavefront”. The wave front spreads laterally but eventually stops (gets pinned) due to depletion of the cytoplasmic substrate, forming stable flat-topped mesa-like concentration profiles. In the absence of spatial cues, wave-pinning models can generate multiple mesas when initiated by random fluctuations (***Mori et al., 2008***). Multiple mesas in the wave-pinning model appear to be “meta-stable” (***Jilkine and Edelstein-Keshet, 2011***; ***Mori et al., 2011***) and do not readily exhibit competition.

An attractive hypothesis for why some cells are uni-polar and others multi-polar would be that these behaviors arise from differences in the biochemical mechanisms of positive feedback, yielding competition in Turing-type or meta-stability in wave-pinning models. However, some Turing-type MCAS models appear to switch to multi-polarity when domain size (***Jilkine and Edelstein-Keshet, 2011***; ***Otsuji et al., 2007***) or protein amount (***Howell et al., 2012***) is increased. Thus, it could be that parameter values (protein concentration, catalytic activity, cell size, etc.) rather than regulatory feedback mechanisms dictate whether uni- and multi-polar outcomes are observed.

Here, we investigate the transient multi-peak scenario, and show that both wave-pinning and Turing-type models are capable of generating uni- or multi-polar outcomes. The switch between uni- and multi-polarity is primarily dictated by a “saturation rule” that is general to MCAS models: Every biologically relevant model has an innate saturation point that sets the maximum local Rho-GTPase concentration. When peaks form such that peak concentrations are well below this saturation point, competition is effective and multi-polar conditions resolve rapidly to a uni-polar steady state. However, if the GTPase concentration in two or more peaks approaches the saturation point, then competition becomes ineffective, and the peaks become meta-stable. Because the saturation rule does not depend on the specifics of the biochemical reactions, our results yield general and testable predictions.

## Results

### MCAS model behaviors

Two-species MCAS systems consist of two partial differential equations (PDEs), governing the dynamics of a slowly diffusing activator (GTP-bound GTPase at the membrane) *u*, and a rapidly diffusing substrate (GDP-bound GTPase in the cytoplasm) *v*. In one spatial dimension, these systems take the general form:

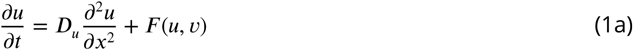

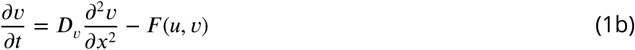

where the dynamics of *u* and *v* are governed by a diffusion term and a reaction term, *F*(*u, v*). To reflect the different compartments (membrane and cytoplasm) populated by the different species, the diffusion constant of *u*, *D_u_*, is typically two orders of magnitude smaller than *D_v_*, so that *u* spreads much more slowly than *v*. *F*(*u, v*) describes the biochemical interconversions between *u* and *v*.

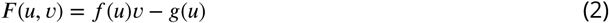

For GTPases, the inactive form of the GTPase *v* is converted to the active form *u* through the action of guanine nucleotide exchange factors (GEFs) *f*(*u*), while *u* is converted to *v* through the action of GTPase activating proteins (GAPs) *g*(*u*). The functions *f*(*u*) and *g*(*u*) take into account potential positive feedback mediated by the active GTPase. Because the inactive GTPase is not thought to participate in biochemical reactions other than as a substrate to produce active GTPase, under the assumption of mass action kinetics *v* appears only in the activation term. As the model assumes only the exchange between *u* and *v*, but not synthesis or degradation of either, the system is mass-conserved, so that the total abundance of the GTPase *M* = *∫*(*u* + *v*)*dx* is a constant over time.

Generation of a GTPase-enriched domain in MCAS models occurs through positive feedback leading to local accumulation of the activator, *u*, and concomitant depletion of the substrate, *v*. Locally depleted *v* is quickly resupplied from the whole cytoplasm due to its high mobility, resulting in a global depletion of *v*. This reduces the net rate, *F*(*u, v*), at which fresh *u* is generated (***Equation 2***), impeding further growth of the *u*-enriched domain, and the system reaches a steady state. At steady state, reaction and diffusion must be balanced at all local positions *x*:

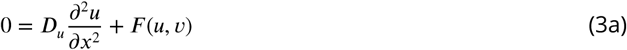

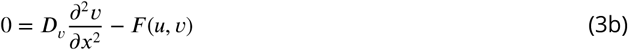

Given a total protein content *M*, these equations govern the steady state peak shape *u*(*x*) and substrate level *v*(*x*) for a single peak in an MCAS model (Further discussed in *Box 1* and Methods section).

Positive feedback can occur through *f*(*u*) (i.e. active GTPase locally stimulates GEF activity) or *g*(*u*) (i.e. active GTPase locally inhibits GAP activity). Examples of feedback via GEF activation include the simple Turing-type model *f*(*u*) = *au*^2^*, g*(*u*) = *bu*, Goryachev’s simplified model *f*(*u*) = *au*^2^ + *cu, g*(*u*) = *bu* (***Goryachev and Pokhilko, 2008***), and Mori’s wave-pinning model 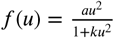, *g*(*u*)=*bu* (***Mori et al., 2008***). Examples of feedback via GAP inhibition include *f*(*u*), = 1, 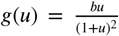, which resembles model I in (***Otsuji et al., 2007***). To illustrate the behaviors of different MCAS models, we simulated examples of Turing-type and wave-pinning MCAS models:

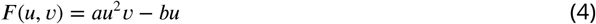

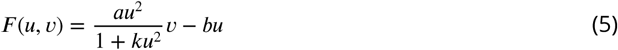

With the appropriate choice of parameters, the Turing-type model (***Equation 4***) yields a peak given any spatial perturbation of the homogeneous steady state, while the wave-pinning model (***Equation 5***) requires a supra-threshold perturbation to destabilize the homogeneous state. The Turing-type model yields a sharp peak at steady state, while the wave-pinning model yields a flat-topped mesa (*Figure 1*E). Simulations with greater total amounts of GTPase *M* yield higher peaks in the Turing-type model, but broader mesas (with the same peak height) in the wave-pinning model (*Figure 1*E), and simulations initiated with two unequal peaks yield rapid competition in the Turing-type model but apparent co-existence in the wave-pinning model (*Figure 1*F). These behaviors are all consistent with previous reports (***Mori et al., 2008***, ***2011***; ***Otsuji et al., 2007***; ***Ozbudak et al., 2005***). To understand these different behaviors, we first revisit the basis for competition.

### Competition between peaks arises from a difference in the ability of unequal peaks to recruit cytoplasmic GTPase

When two unequal peaks are present in the same domain, each peak recruits GTPase from the cytoplasm, thereby globally depleting cytoplasmic GTPase. As exchange of GTPase between each peak and the cytoplasm is dynamic, the two peaks are effectively recruiting GTPase from one another. If the larger peak (the one that contains more GTPase) recruits GTPase more effectively, it will grow at the expense of the smaller peak, eventually yielding a uni-polar outcome (*Figure 2*A, scenario 1). If instead, the smaller peak recruits GTPase more effectively, then it will grow while the larger peak shrinks, eventually yielding two equal peaks, as observed in some more complex models (***Howell et al., 2012***) (*Figure 2*A, scenario 2). If two unequal peaks recruit GTPase equally, then the two unequal peaks would simply coexist (*Figure 2*A, scenario 3).

**Figure 2.**
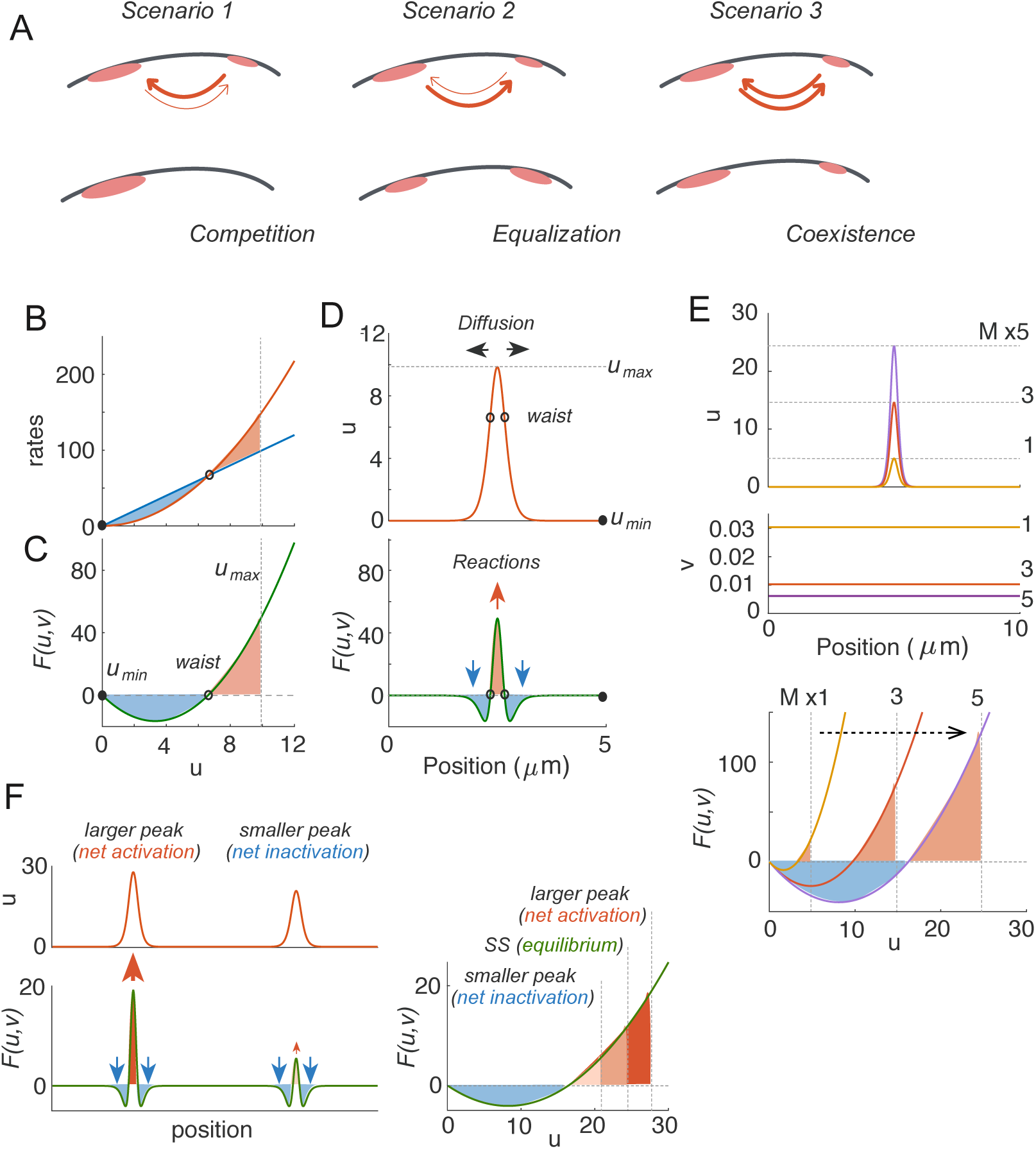
The basis for competition. A) Possible outcomes when there are two unequal clusters of Rho-GTPase in the same cell. Scenario 1: competition occurs if larger clusters recruit GTPase more efficiently than smaller clusters. Scenario 2: equalization occurs if smaller clusters recruit GTPase more efficiently than larger clusters. Scenario 3: co-existence occurs if both clusters recruit GTPase equally well. B-F: Turing-type model with *D_v_* → ∞. B) Rate balance plot: activation and inactivation rates are balanced at two fixed points of *F* (*u, v*). Filled circle indicates stable fixed point, and empty circle indicates unstable fixed point. C) Net activation (shaded red) and net inactivation (shaded blue) from the trough (*u*_min_) to the top (*u*_max_) of the peak must be balanced at steady state (*Box 1*). This determines the peak height (*u*_max_). D) Net activation at the center of the peak is balanced by diffusion, which drives GTPase towards the flanks, where there is net inactivation. E) If total GTPase content is raised, the model generates higher peaks (larger *u*_max_), accompanied by more severely depleted *v*, which lowers *F*(*u, v*) such that the blue and red shaded areas are once again balanced. F) When two peaks are present, they share the same *v* and hence the same *F*(*u, v*) curve. The larger peak will always have excess net activation, and the smaller peak will always have excess net inactivation, so competition is inevitable. Parameter values used: *a* = 1 *μm*^2^, *b* = 1 *s*^−1^ and *D_u_* = 0.01 *μm*^2^*s*^−1^, *D_v_* = ∞. All models were simulated on domain size *L* = 10 *μm*.

To understand how these considerations play out for different peaks, we need to know which peak will recruit more GTPase. To assess how much GTPase would be recruited to a specific peak, consider first the Turing-type model (***Equation 4***) in the limit *D_v_* → ∞. This model combines a quadratic (in *u*) activation term with a linear inactivation term (*Figure 2*B). Thus, for a fixed value of *v*, there are two values of *u* at which activation and inactivation balance each other precisely (i.e. fixed points of the net activation curve *F*(*u, v*), *Figure 2*C). Given the concentration profile of a peak (*Figure 2*D, upper panel), *F*(*u, v*) determines whether any given location on the membrane will gain GTPase from the cytoplasm or lose GTPase to the cytoplasm (*Figure 2*D, lower panel). At the trough in *Figure 2*D (*u*_min_), *u* approaches the lower fixed point of *F*(*u, v*), yielding no net gain or loss of GTPase. On the lower flanks of the peak, *u* values lie between the two fixed points, and inactivation outpaces activation, so there is a net loss of *u* (*Figure 2*B, C and D). When *u* rises above the higher fixed point of *F*(*u, v*), up until the top of the peak (*u*_max_), there is net recruitment of GTPase from the cytoplasm (*Figure 2*B, C, and D). At steady state, the net loss from *u*_min_ to the higher fixed point (blue area in *Figure 2*B, C) is balanced by the net recruitment from the higher fixed point to *u*_max_ (red area in *Figure 2*B, C, *Box 1*). Additionally, diffusion from the center of the peak to the flanks balances these flows of GTPase, requiring a sharp-topped peak (where negative 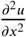 counteracts net recruitment at the center: ***Equation 3***) (*Figure 2*D). We could generate a larger peak by increasing the total GTPase content (*M*) of the system: positive feedback would then drive more GTPase into the peak, so *u*_max_ would increase to yield greater net activation in the center of the peak, resulting in more severe depletion of cytoplasmic *v* and hence shifting *F*(*u, v*) (*Figure 2*E). At steady state, the red and the blue areas (though each larger than for the smaller peak) would once again be equal.

Now consider a scenario in which two unequal peaks are present in the same domain. Both peaks would grow until cytoplasmic *v* becomes sufficiently depleted. In the limit where *D_v_* → ∞, *v* will reach this concentration, *v*^*^, throughout the cytoplasm shared by both peaks. Therefore, the same net reaction curve will apply to both peaks, but they will have a different *u_max_* (*Figure 2*F). The overall recruitment or loss of GTPase for each peak *u*(*x*) sharing a common *v*^*^ is given by:

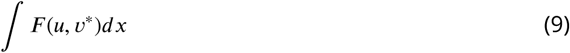

#### Box 1. Intuitive description of the steady state solutions by analogy to Newtonian physics.

A steady state is reached when the three fluxes, diffusion, activation (conversion of *v* to *u*) and inactivation (conversion of *u* to *v*), reach equilibrium at all spatial positions (***Equation 3***). To satisfy this condition, the total difference between the activation and inactivation curves (area shaded from the bottom to the top of the peak in *Figure 2*B-E) is zero. This can be understood as follows.

First, at steady state, we can rewrite ***Equation 3a*** in the form

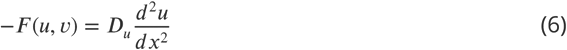

In the general scenario *D_v_ <* ∞, *v* = −*D_u_*/*D_v_* +*q* (***Equation 11***), where *q* is a constant representing the basal substrate level. In the limit *D_v_* → ∞, *v* = *q*.

This equation (***Equation 6***) has the same form as Newton’s second law *F* = *ma*, where −*F*(*u, q*) is analogous to the force, *D_u_* is analogous to the mass, *u* is analogous to the position, and “*x*” is analogous to time. The relationship between the “force” and the “position” (*u*) is governed by −*F*(*u, q*). Equivalent to this equation is the condition for conservation of energy,

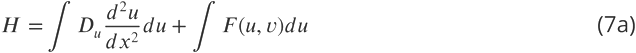

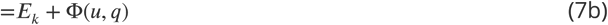

where the constant H is the total energy and the kinetic and potential energies are defined by the relations

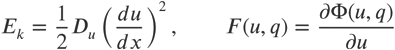

respectively.

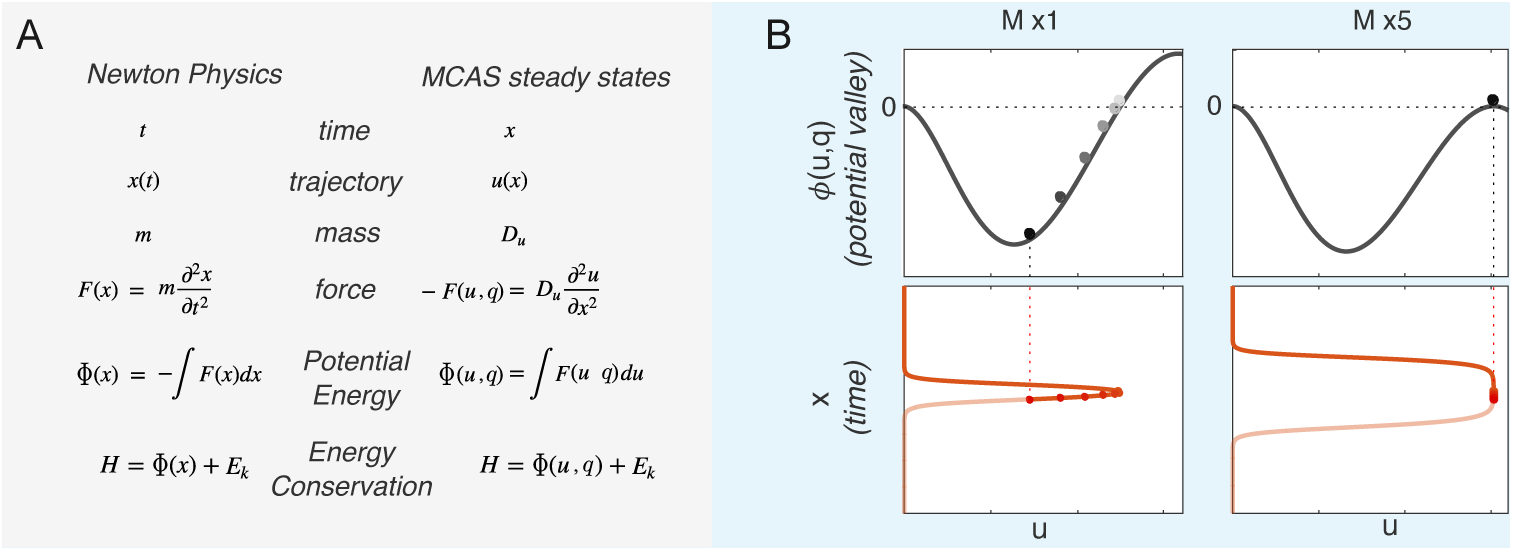

The **concentration profile** *u*(*x*) is now analogous to the **spatiotemporal trajectory** of a ball rolling under gravity on a 1D potential valley defined by Φ(*u, q*) (B, left panel; Video 1). When the ball is at the left edge of the valley, which corresponds to the bottom of a concentration profile (*u*_min_), it has maximum potential energy and zero kinetic energy. As the ball rolls down, potential energy transforms into kinetic energy, until it reaches the bottom of the valley, where it has maximal kinetic energy and minimal potential energy. This corresponds to the waist of the peak in the concentration profile, where the slope of *u*(*du*/*dx*) is steepest. The ball slows down as it rolls up the other side of the valley, and eventually stops at a position analogous to the top of the concentration peak(*u*_max_).

Due to energy conservation, the ball must reach the same level at the right edge as at the left edge of the valley. Since Φ is the integral of *F*(*u, v*), energy conservation in the Newtonian analogy demands that the area between the activation curve and the inactivation curve from the bottom to the top of the peak sums up to zero. This has been referred to as the “**wave-pinning condition**” (***Mori et al., 2008***), but applies to other MCAS models as well.

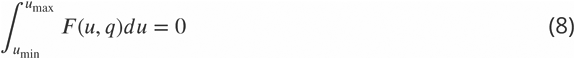

With the Newtonian analogy, it follows that the flat top of a mesa corresponds to the ball staying at *u*_max_ for a long “time” *x*. This only occurs when the ball has just enough energy to reach the top of the potential valley (local maximum of Φ) on the right, where the force on the ball (*F*(*u, q*)) approaches zero (B, right panel; Video 2). Therefore, the conditions for a mesa to occur are 1) that *F*(*u, q*) has a third fixed point, and 2) that the substrate level *q* is low enough so the top of the potential valley (local maximum of Φ) reaches the same potential as the left edge of the valley, which means that *u*_max_ approaches the third fixed point of *F*(*u, q*).

The higher the *u*_max_ of a peak, the larger the overall recruitment, demonstrating that the larger peak has a stronger “recruitment power” than the smaller peak (*Figure 2*F). Thus, in a scenario with unequal peaks in the same domain, the larger peak experiences a net gain of GTPase, while the smaller peak experiences a net loss, further exacerbating the inequality between the two peaks until the smaller peak is eliminated. The Turing model (***Equation 4***) with *D_v_* → ∞ always competes to yield a uni-polar endpoint (scenario 1 in *Figure 2*A).

The argument above requires only mass-conservation and non-linear positive feedback, which is a core requirement for polarization in general (***Gierer and Meinhardt, 1972***). Therefore, it would seem that all MCAS models should compete, regardless of the specific *F*(*u, v*). To verify this, we generated steady states with two symmetric peaks in a domain, and performed linear stability analysis to show that such steady states are unstable (See Methods section, *Figure 9*). Perturbations that destabilize the steady state yield either competition between the peaks or merging of the peaks. Here we focus on competition. Our analysis in the limit of *D_v_* → ∞ indicates that given sufficient time, two peaks will always compete to produce a single peak. This result does not depend on the form of *F*(*u, v*).

### Competition slows down dramatically due to saturation

If competition (scenario 1 in *Figure 2*A) applies to all MCAS models, then why did we not observe competition in simulations of the Wave-pinning model (*Figure 1*G)? In contrast to the Turing-type model (***Equation 4***), the reaction term of the Wave-pinning model (***Equation 5***) has saturable positive feedback, introducing a third fixed point in *F*(*u, v*) (*Figure 3*A). When the total protein content in the system is small, *u*_max_ does not approach this fixed point (*Figure 3*B). Under these conditions, sharp-topped peaks compete with each other to yield a uni-polar outcome, as with the Turing-type model (*Figure 3*C). But when protein content of the peak is increased, *u*_max_ approaches the third fixed point, and the net activation rate *F*(*u, v*) at the top of the peak approaches zero (*Figure 3*B). To satisfy the steady-state condition (***Equation 3a***), 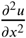 must also approach zero. In other words, the top of the peak must broaden to become a flat-topped mesa. Once this occurs, increasing *M* only negligibly increases *u*_max_, and instead of developing higher peaks the model develops broader mesas with comparable *u*_max_ (*Figure 3*B). We shall call this maximum value the “saturation point” (*u*_sat_) of the model.

**Figure 3.**
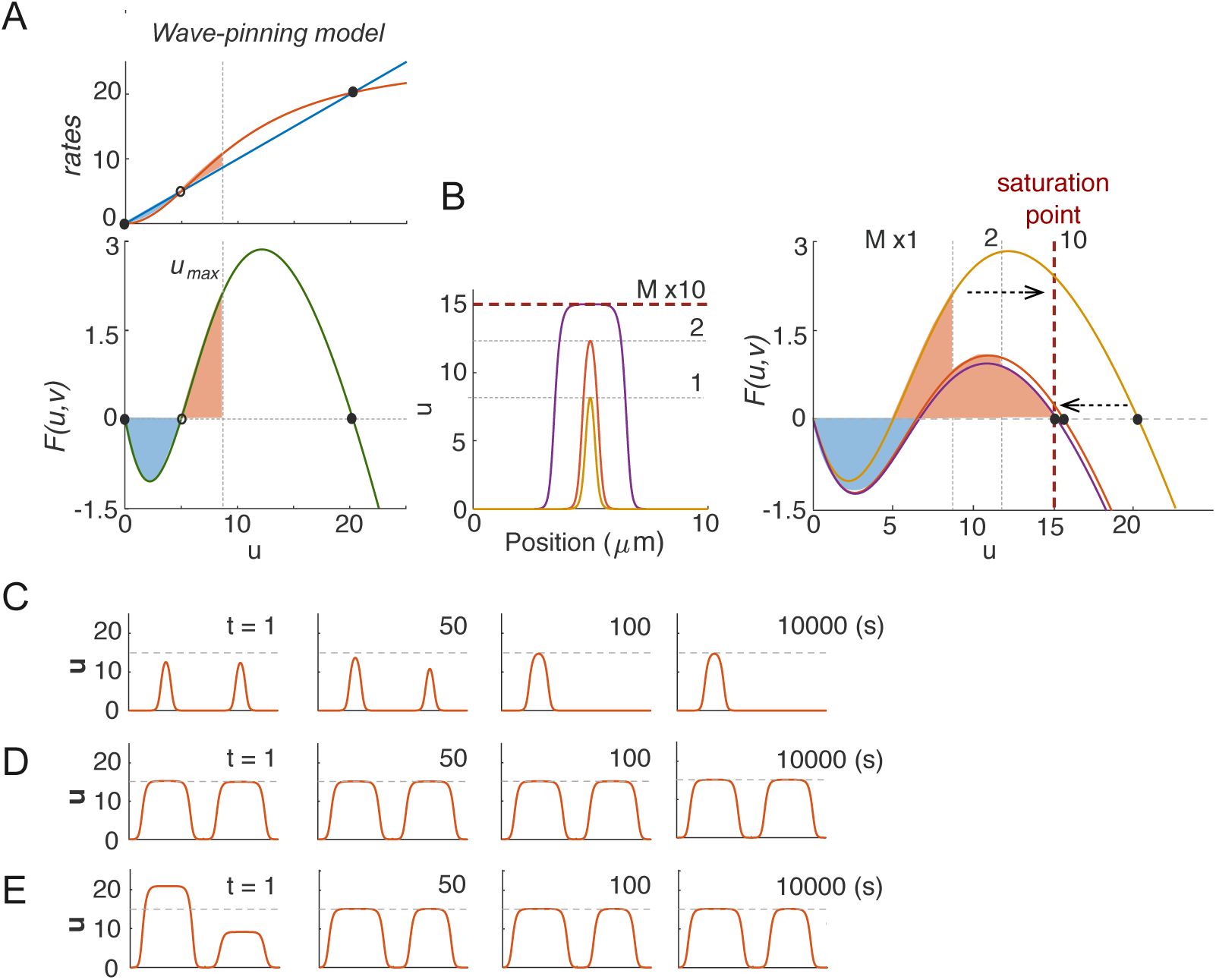
Competition in Wave-pinning models. A) The wave-pinning model has a saturable activation term, introducing a third fixed point in *F*(*u, v*). Dashed line indicates *u*_sat_. Circles indicate stable (filled) and unstable (empty) fixed points. B) As total GTPase levels *M* increase, the peaks get higher until *u*_max_ reaches the saturation point (the third fixed point), after which peaks broaden into mesas. C) With *M* = 40, two identical peaks were perturbed by 1% at *t* = 0 *s*. The resulting competition led to a single-peak steady state within 100 *s*. D) With *M* = 200, the same 1% perturbation did not result in noticeable competition in 10000 *s*. E) Starting from the same two-peak steady state as in D, we introduced a large 50% perturbation. The two mesas quickly evolved back to the original *u*_max_, and then persisted for 10000 *s*. *k* = 0.01 *μm*^2^. Other parameters same as *Figure 2*.

When *u*_max_ approached the saturation point *u*_sat_, simulations with two flat-topped peaks did not show obvious competition (*Figure 3*D). Applying a drastic perturbation in which 50% of the GTPase in one peak was transferred to the other led to a rapid adjustment with both peaks returning to an almost identical *u*_max_ but with different peak widths, after which the unequal peaks co-existed for prolonged simulation times (*Figure 3*E) (Note that the two peaks did not “equalize”: they retained unequal total GTPase content.) Thus, the same model can yield rapid competition or competition so slow as to yield prolonged co-existence, simply as a result of varying the total amount of GTPase in the system.

To investigate more broadly how model parameters might influence the timescale of competition between peaks, we simulated competition between two unequal peaks in the Wave-pinning model, in the limit with *D_v_* → ∞. If we start with a two-peak steady state and noise, the two peaks will eventually resolve to one, given sufficient time. As a measure of competition time that should be insensitive to the precise degree of the noise, we tracked the time it took for unequal peaks with active GTPase content ratio 3:2 to evolve to a content ratio of 99:1. Parameter changes caused dramatic changes in competition times, color coded on a log scale in *Figure 4*A. Notably, increasing *M* always led to slower competition (*Figure 4*A, left panel). As discussed above, increasing GTPase content causes *u*_max_ to approach the saturation point. Defining a saturation index in terms of how closely *u*_max_ at the two-peak steady state approached the saturation point ((*u*_sat_ − *u*_max_)/*u*_sat_), we found that the effects of varying parameters on the saturation index closely paralleled the parameter effects on the timescale of competition (*Figure 4*A, right panel). A similar congruence was observed using peak width as a different measure of how closely the system approaches saturation (*Figure 4*-Figure supplement 1). These findings suggest that a large majority of the variation in competition times can be explained simply by the degree to which peaks in the model approach the saturation point.

**Figure 4.**
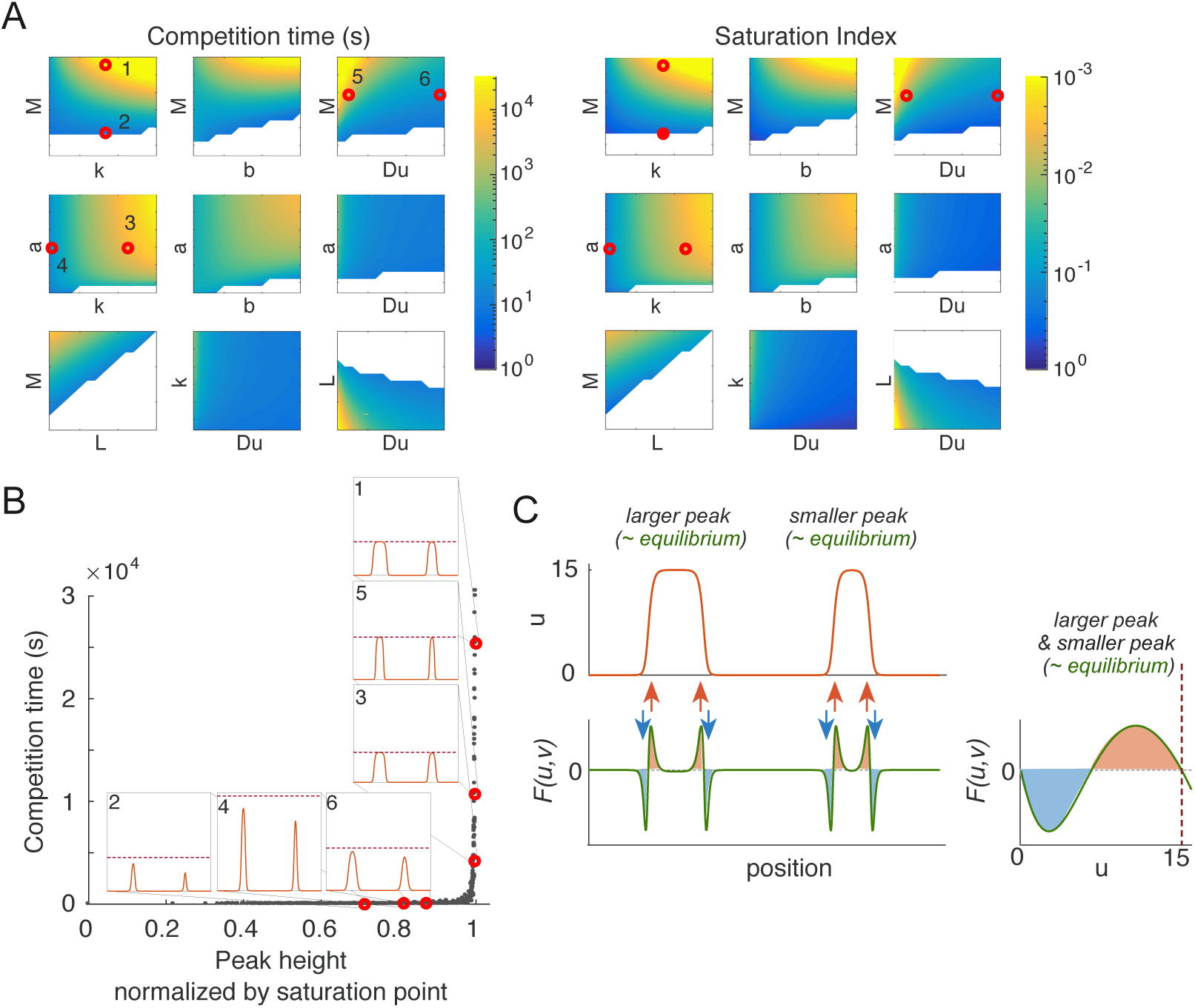
Saturation is a major contributor to differences in competition times. A) Competition time and saturation are tightly correlated. Competition time (*s*) is shown in color (note log scale). Saturation index is defined here as (*u*_sat_ − *u*_max_)/*u*_sat_, and colored in inverse log scale (smaller saturation index indicates peaks are closer to saturation). Basal parameters: *a* = 1 *μm*^2^*s*^−1^, *b* = 1 *s*^−1^, *k* = 0.01 *μm*^2^ and *D_u_* = 0.01 *μm*^2^*s*^−1^, *D_v_* = ∞, *M* = 40, *L* = 20 *μm*. Each color plot represents a 15-fold parameter variation from 0.2x ~3x of the basal parameter value. White regions indicate parts of parameter space where polarized states collapse to homogeneous states. Numbered red dots correspond to the simulations illustrated in the inset of panel B). B) Each of the simulations performed for panel A) is plotted as one dot. Competition time (Y axis) is plotted against peak height *u*_max_ normalized to the saturation point *u*_sat_ for that simulation (X axis). Inset graphs indicate starting conditions for the selected simulations with parameters indicated by red dots in A). C) When two mesas coexist, they share the same *F*(*u*, *v*) curve and almost the same *u*_max_. Thus, the wider peak has a negligible recruitment advantage over the narrower one. **Figure 4-Figure supplement 1.** Peak width, *ℓ*_mesa_, is a robust indicator of saturation over a broad range of system parameters. Data points collected from simulations in *Figure 4*A.

If we plot competition time against *u*_max_ normalized to the saturation point, all of the simulations with different parameter values display one of two clearly distinct behaviors (*Figure 4*B). Parameter changes can alter GTPase content in the peaks (*Figure 4*A,B, point 1 vs 2), the saturation point (point 3 vs 4), or the shapes of the peaks (point 5 vs 6). In all cases, whenever *u*_max_ is not close to saturation, competition occurs rapidly. Conversely, as *u*_max_ approaches the saturation point, competition slows sharply and the two-peak situation becomes meta-stable, resembling the co-existence scenario 3 in *Figure 2*A.

The basis for the drastically slowed competition in simulations with peaks close to saturation can be intuitively understood in terms of each peak’s “recruitment power” (***Equation 9***). When peaks approach saturation, unequal peaks differ in width but have almost identical *u*_max_ and hence only a negligible difference in recruitment power (*Figure 4*C). At the flat tops of the peaks, *F*(*u, v*) = 0, so the peak tops (of any width) do not contribute to overall recruitment. For that reason, the extra GTPase in a broader peak does not give it a significant advantage over the narrower peak, and the driving force for competition is negligible.

Analysis of the eigenvalues from linear stability analysis of this system shows that the timescale of competition slows exponentially as the peaks increase in width. This conclusion, again, is general to all MCAS models and can be applied to all formulations *F*(*u, v*) that allow a third fixed point (See methods section, *Figure 9*).

### Local cytoplasmic depletion also leads to saturation and slow competition

When cytoplasmic diffusion is finite (*D_v_ <* ∞), a saturation point emerges even if there is no explicit saturation in the reaction term. With finite *D_v_*, increasing *M* in the Turing-type model (***Equation 4***) yields flat-topped peaks that become broader as *M* increases (*Figure 5*A), similar to that seen with the wave-pinning model (***Equation 5***).

**Figure 5.**
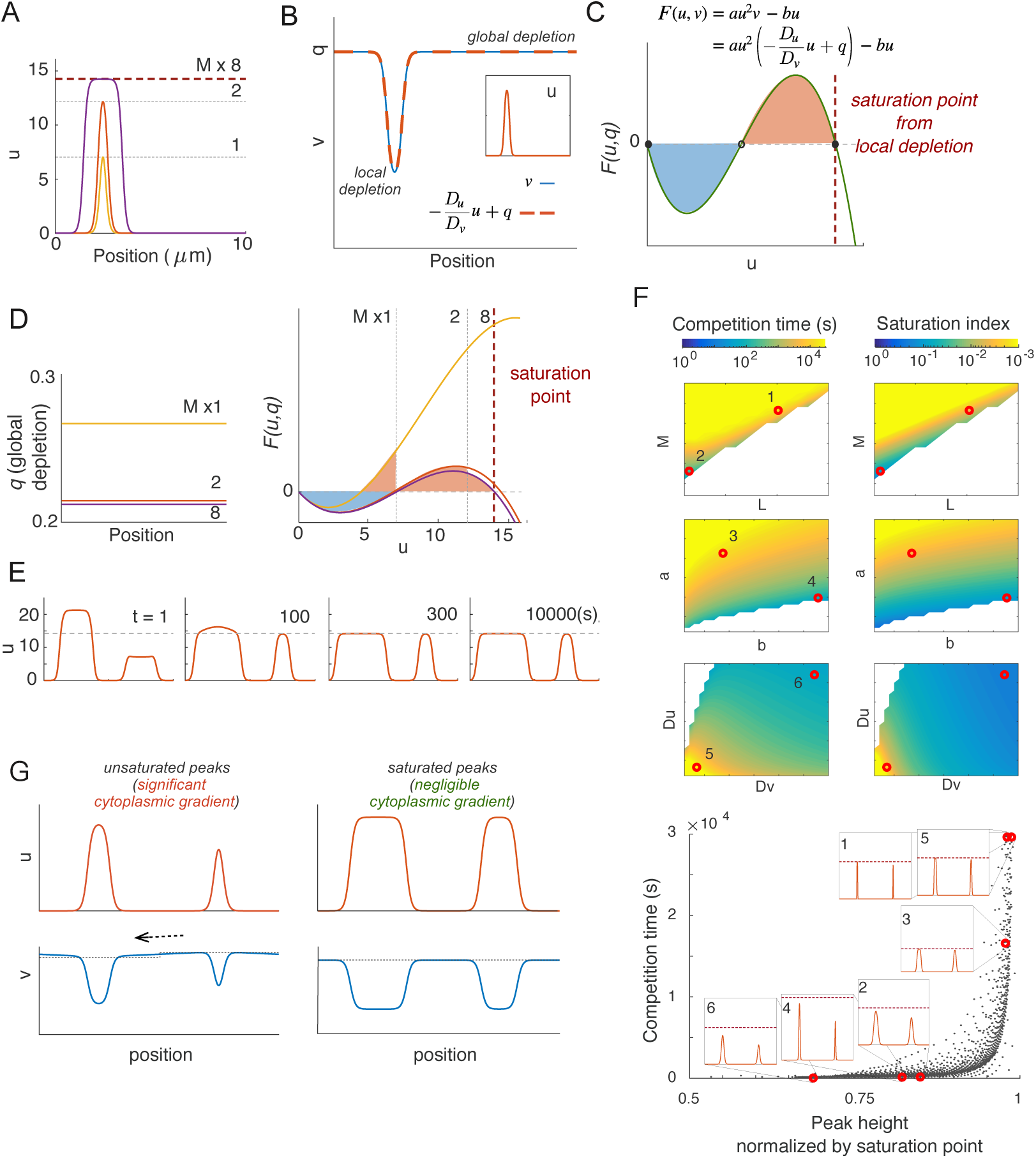
Local substrate depletion leads to saturation and slow competition. A) Turing-type model with *D_v_ <* ∞ displays a transition between sharp peaks and flat-topped mesas with increased *M*. B) Local depletion of *v* in the cytoplasm beneath the peak results in a linear relationship between the concentration profile of *v* and *u*. Inset indicates *u* profile. C) The effect of local depletion transforms the reaction term of the Turing-type model from a quadratic *F*(*u*, *v*) to a cubic *F*(*u, q*), yielding a third fixed point. D) The cubic reaction term *F*(*u, q*) results in a behavior similar to that of the wave-pinning model: When *M* is low, *q* is high, and the peak is sharp; when *M* increases, depletion of cytoplasmic substrate makes *F*(*u, q*) drop, and *u*_max_ eventually approaches a saturation point. E) Peaks saturated by local depletion are meta-stable. F) Saturation index correlates with competition timescale. Simulations and display as in *Figure* *4*A,B. Parameter variations in *a* vs *b* and *D_u_* vs *D_v_* consist of 30x30 simulations each of 0.1x ~3x of the basal parameter values. Parameter variations in *M* vs *L* consists of 15x15 simulations of 0.2 ~3x basal parameter values. Basal parameters are as in *Figure* *4*A, except that *D_v_* = 1 *μm*^2^*s*^−1^. Graph shows all simulations plotted as in *Figure* *4*B, with illustrative simulations corresponding to numbered red dots. G) When *D_v_*, the basal cytoplasmic substrate concentration underneath each peak (shown in dashed lines) quickly reaches a quasi-steady state with the peak. The stronger the recruitment power of the peak, the lower the basal cytoplasmic substrate level. This creates a cytoplasmic gradient when two peaks have different recruitment power, resulting in a cytoplasmic flux towards the larger peak. The gradient becomes negligible when both peaks are saturated, resulting in meta-stable peaks.

To understand this behavior, recall that at steady state, ***Equation 3*** must hold. Adding ***Equation 3a*** and ***Equation 3b***, integrating and enforcing the periodic boundary condition yields a linear relationship between *u* and *v*, regardless of the reaction term:

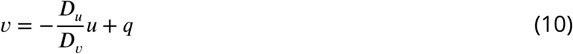

where *q* is a constant over space that represents the baseline cytoplasmic level of *v*, due to global depletion of substrate GTPase. This reflects the fact that in addition to global substrate depletion, activation due to positive feedback depletes *v* locally under a peak of *u*, creating a “dip” in the concentration of the cytoplasmic GTPase *v* that corresponds to the peak of *u* in a linear manner (*Figure 5*B).

Local depletion results in an emergent saturation effect, because substituting ***Equation 10*** into the reaction term of the Turing type model (***Equation 4***) gives:

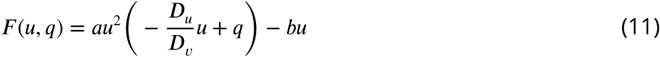

This new reaction term *F*(*u, q*) is a cubic in *u*, and can have three fixed points (*Figure 5*C). The upper fixed point reflects the *u* concentration at which local depletion of *v* precisely balances the net recruitment of *u*, yielding an emergent saturation point. Thus, even when there is no saturation inherent in the reaction term of the model, local depletion of *v* under the peak acts to limit local production of *u*, introducing a saturation effect. Given sufficient total mass *M*, *u*_max_ approaches this saturation point, resulting in a flat-topped peak for reasons described above with the wave-pinning model (*Figure 5*D). In this case, it is possible to derive a simple expression for the saturation point:

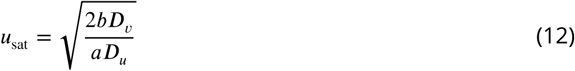

As with saturation due to the wave-pinning reaction term, saturation by local depletion also slowed competition dramatically, leading to meta-stable peaks (*Figure 5*E). Exploration of a wide parameter range indicated that as with saturation via the reaction term, saturation due to local depletion of substrate is also a dominant contributor to the timescale of competition (*Figure 5*F).

When *D_v_ <* ∞, two unequal peaks no longer “see” the same level of substrate, *v*. Instead, the local *v* rapidly reaches a quasi steady-state with each peak (*Figure 5*G). When two unsaturated peaks coexist, the higher peak has a stronger recruitment power for reasons discussed in *Figure 2*F. This drives a greater depletion and hence lower baseline of *v* under the higher peak, generating a cytoplasmic *v* gradient that drives a flow of GTPase towards the higher peak, and hence competition (*Figure 5*G). In contrast, when two unequal but saturated peaks coexist, they have similar recruitment power, so there is a negligible cytoplasmic gradient, and competition occurs on a dramatically slower timescale.

### Unifying Turing and Wave-pinning models

As the Turing-type and Wave-pinning models behaved similarly with regard to to competition and saturation, we revisited their behavior with regard to diffusion-driven instability and wave-like spread. We first explored the behavior of simulations of the Wave-pinning (***Equation 5***) and Turing-type (***Equation 4***) models starting from the homogeneous steady state with random noise for *u*. In both cases, multiple peaks formed and then rapidly competed. As the winning peaks grew to approach the saturation point, competition slowed dramatically (*Figure 6*A). Linear stability analysis of the wave-pinning model (see Methods section) confirmed that there is a parameter regime in which this model is Turing unstable (*Figure 6*C).

**Figure 6.**
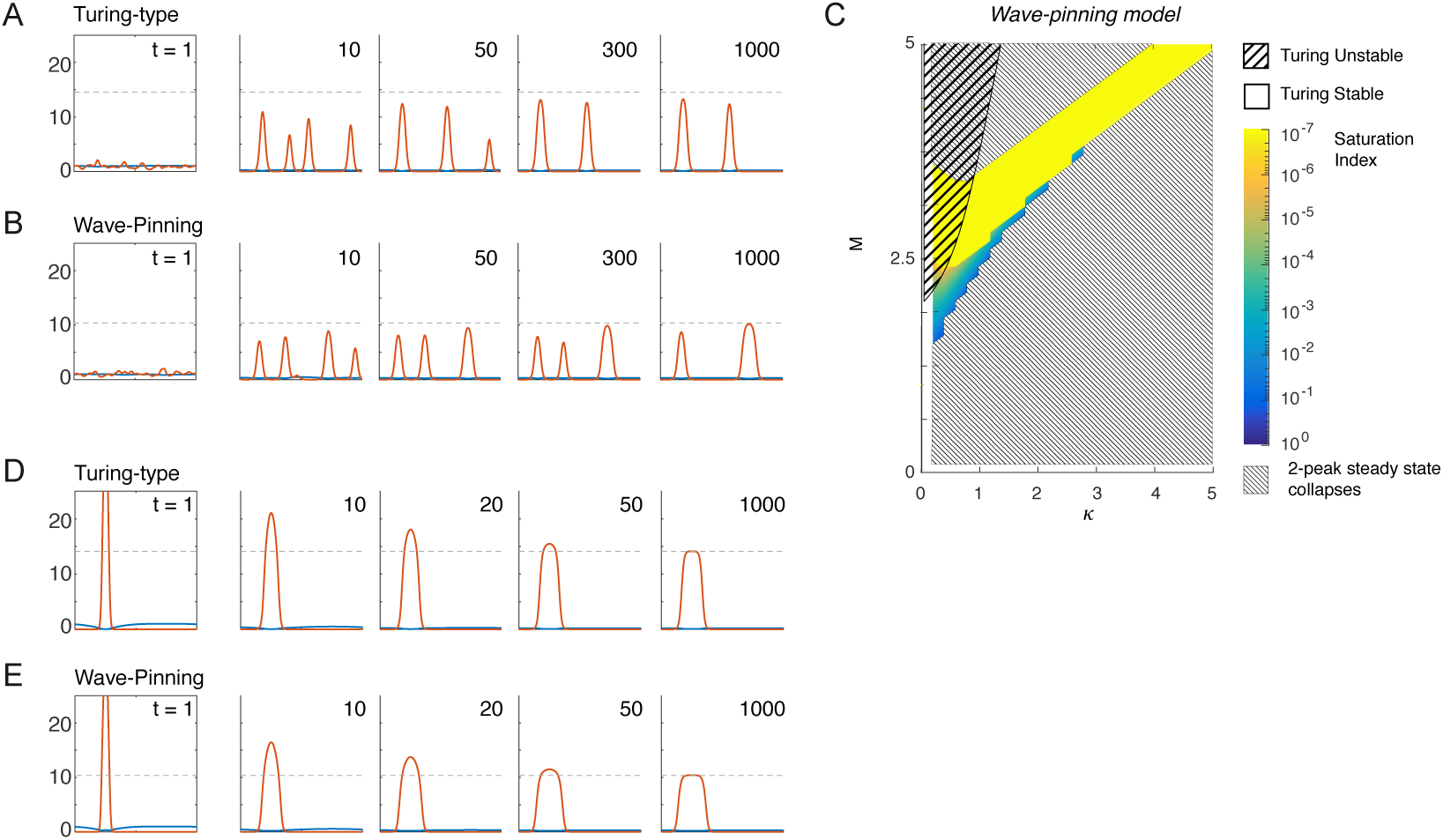
Turing instability and wave-pinning behavior in both the Turing-type and the Wave-Pinning models. A, B) Simulations of Turing-type (***Equation 4***) and Wave-Pinning model (*Equation 5*). *u* is indicated in red, *v* in blue. Dashed line indicates the saturation point. *M* = 2, *D_u_* = 0.01 *μm*^2^*s*^−1^, *D_v_* = 1 *μm*^2^*s*^−1^ for both models; *k* = 0 *μm*^2^ for the Turing-type model and *k* = 0.0 *μm*^2^ for the Wave-Pinning model. C) Behaviors of the Wave-Pinning model. Stability of the homogeneous steady state was calculated as in methods section. Saturation index was extracted from simulated 2-peak steady states, plotted in color in log scale. Uncolored regions indicate parameter spaces where the 2-peak steady state collapses to the homogeneous steady state. D,E) Both the Turing and the Wave-Pinning model can manifest wave-pinning behavior. Initial conditions for both simulations are *u* = 0, *v* = *M*, with a spike in *u* that triggers the wave. *u* is indicated in red, *v* in blue. Parameter values are the same as in A and B, except that *M* = 2.6.

Wave-pinning dynamics are thought to depend on a bi-stable system (***Mori et al., 2008***, ***2011***). As we showed that Turing-type models can exhibit bi-stability due to local depletion of cytoplasmic substrate, they too should be able to manifest wave-pinning dynamics. Indeed, if we start a simulation with one large spike of *u* and all other material as *v*, the spike triggers positive feedback and expands in a wave-like manner. As the wave spreads, *v* is depleted, until eventually the wave-pinning condition ***Equation 8*** is satisfied, and the wave stops when the top of the peak corresponds to the saturation point *u*_sat_ of each model. This behavior is seen in both Turing-type and Wave-pinning models without discernible qualitative differences.

In summary, MCAS models may exhibit Turing instability or Wave-pinning dynamics, and may compete effectively or co-exist, depending on parameters. The “typical” behaviors of Turing and Wave-pinning models simply represent behaviors of MCAS models in specific parameter subspaces. This view is consistent with a recent review (***Goryachev and Leda, 2017***).

### Effect of increasing distance between peaks

During competition, GTPase is transferred from the “losing” peak to the “winning” peak through the cytoplasm. Thus, increased distance between the peaks or a decreased diffusion constant in the cytoplasm would be expected to slow the transfer and hence slow competition (an effect not seen when *D_v_* → ∞). To assess how effective increased distance could be in slowing competition, we initially considered the effect of increasing cell size while keeping overall GTPase concentration constant (*Figure 7*, gray line). Competition slowed dramatically as domain size *L* was increased, but this does not distinguish whether increasing distance between peaks or increasing total GTPase content *M* (moving the peaks closer to saturation) is responsible for the slowing. Increasing *L* without changing *M* resulted in GTPase dilution and hence smaller peaks that competed more rapidly despite the increased distance between peaks (*Figure 7*, blue line). To maintain equivalent peaks, we increased *L* while adding the exact amount of GTPase required to fill the cytoplasm in the extended domain so that the amount of GTPase *in the peaks* remained constant. This scenario allowed us to quantify the effect of increasing distance between peaks without confounding changes in peak size. The result was that competition became slower in a sub-linear manner with distance (*Figure 7*, red line). Thus, distance between peaks can slow competition, but does so in a much more gradual manner than the approach to saturation.

**Figure 7.**
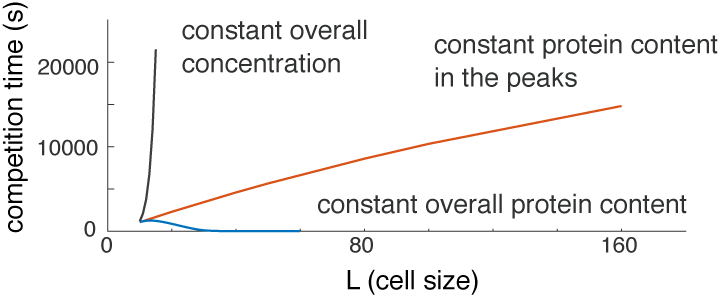
Effect of domain size on competition time. Effect of expanding the domain size on competition time. Gray: overall concentration was set to constant as L increases (proportional increase of total protein content in the system *M*; peaks saturate). Blue: overall protein content constant (peaks shrink to feed the larger cytoplasm). Red: protein content in the peaks is maintained constant (identical peak shape).

### Other MCAS models also link competition timescale to saturation

Our analysis has focused on specific illustrative models, but many other forms of *F*(*u, v*) in ***Equation 2*** can also support polarization. For example, positive feedback strength may vary, yielding different exponents for the activation term (e.g. *f*(*u*) = *u*^1.2^ with weak feedback, or *f*(*u*) = *u*^3^ with strong feedback). Or, positive feedback may operate by reducing inactivation rather than by increasing activation (e.g. *f*(*u*) = 1*,g*(*u*) = *u*/(1 + *u*^2^)). Or, positive feedback may be accompanied by negative feedback, as proposed for the yeast polarity circuit (***Howell et al., 2012***; ***Kuo et al., 2014***) (e.g. *f*(*u*) = *u*^2^ − *cu*^4^). As local cytoplasmic depletion is a universal mechanism of saturation, we would expect that competition time slows down as the system approaches saturation in all of these models. Indeed, all of these variations displayed a saturation point, leading to a transition from sharp peaks to mesas as *M* was increased. And in each case, the change in peak shape was accompanied by a dramatic slowing of competition (*Figure 8*A-E). This suggests that our findings are broadly applicable to MCAS models.

**Figure 8.**
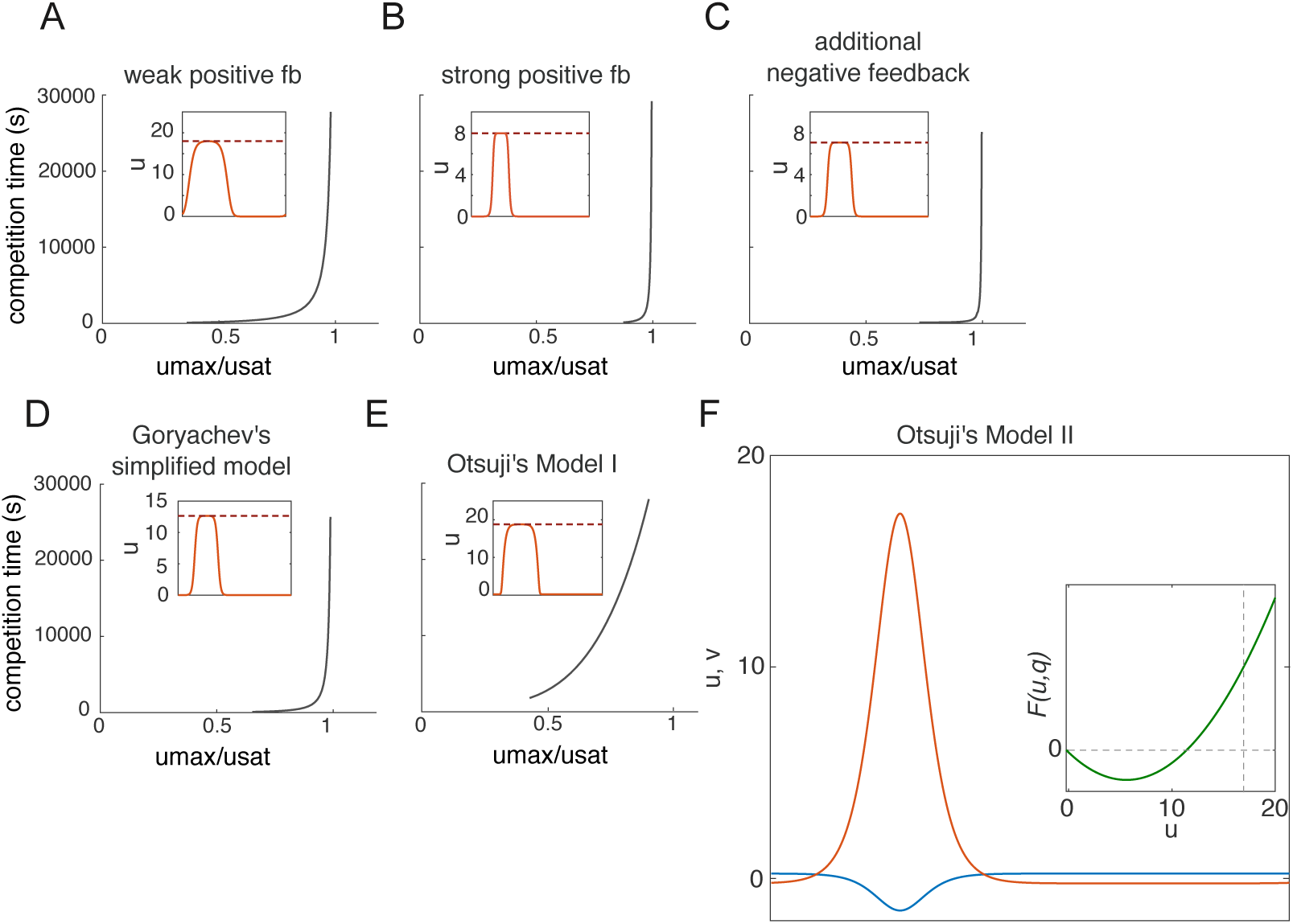
Other MCAS models also link competition timescale to saturation. Competition time as a function of saturation for other MCAS models. Insets: peak shape upon reaching saturation. Red dashed lines indicate saturation point. A) weak positive feedback, *F*(*u*, *v*) = *u*^1.2^*v* − *u*; B) strong positive feedback, *F*(*u*, *v*) = *u*^3^*v* − *u*; C) additional negative feedback *F*(*u*, *v*) = (*u*^2^ − 0.01*u*^4^)*v* − *u*; D) Goryachev’s simplified model *F*(*u*, *v*) = (*u*^2^ + *u*)*v* − *u* (***Goryachev and Pokhilko, 2008***); E) Otsuji’s model 1 *F*(*u*, *v*) = *a*_1_*v* − *a*_1_(*u* + *v*)/[*a*_2_(*u* + *v*) + 1]^2^ with the original parameters described in (***Otsuji et al., 2007***). In each instance, competition time slows down dramatically as peaks saturate. F) In Otsuji's model 2 with the original parameters, *F*(*u*, *v*) = *a*_1_(*u* + *v*)[(*D_u_*/*D_v_u* + *v*)(*u* + *v*) − *a*_2_] (***Otsuji et al., 2007***), saturation is avoided by allowing negative values of *u* or *v*.

The only counterexample we have encountered so far is model II from ***Otsuji et al.*** (***2007***), where

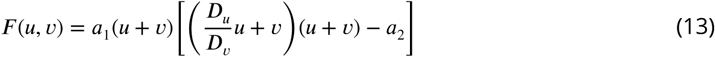

Unlike other reaction terms based on mass action kinetics (***Equation 2***), this reaction term is not dependent on *v*, but rather on the combined concentration of *u* and *v*. Thus, activation in this model is no longer restricted by *v* depletion as in the other models mentioned above, and *v* can assume negative values when *u* is high, avoiding saturation (*Figure 8*F). This eliminates the effect of local depletion: When *v* is substituted with 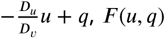 is a curve lacking a third fixed point (and hence lacking saturation). However, as concentrations of *u* or *v* cannot be negative in cells, this model is not physiologically relevant.

## Discussion

### Competition between peaks obeys a “saturation rule”

Early studies on MCAS systems emphasized that different models can display different behaviors, including Turing instability and wave-pinning dynamics. In the parameter regimes examined, Turing-type model peaks displayed rapid competition, while wave-pinning model peaks co-existed, suggesting that competition might be linked to model architecture. However, our findings do not support a categorical distinction between model types. Indeed, a classical wave-pinning model can exhibit Turing instability depending on parameters, and a classical Turing-type model can exhibit wave-pinning behavior when the total GTPase amount is increased (*Figure 6*) (***Goryachev and Leda, 2017***).

Instead of categorizing MCAS models into different types, our findings lead us to propose that the dynamics of competition between peaks obey a “saturation rule”. We suggest that competition between activator peaks for the shared pool of cytoplasmic substrate (scenario 1 in *Figure 2*A) is universal for all biologically relevant MCAS models. However, each model encodes a calculable, parameter-dependent saturation point, such that the peak activator concentration cannot exceed that level at a polarized steady state. As the peak activator concentration approaches the saturation point, the difference between unequal peaks in terms of their ability to recruit cytoplasmic substrates becomes negligible, leading to dramatically slower competition and effective co-existence between peaks (scenario 3 in *Figure 2*A). Varying parameters affects competition time predominantly by affecting the degree to which competing peaks approach the saturation point.

### Biological implications of the saturation rule

The models considered in this report represent a drastically simplified system compared to any biological system. Three simplifying assumptions are particularly noteworthy. First, we considered only a single spatial dimension, whereas cell membranes are two-dimensional and the cytoplasm is three-dimensional. Higher dimensions introduce additional factors such as curvature (***Mori et al., 2011***; ***Ramirez et al., 2015***) that may also affect competition. Second, because polarization phenomena often employ stable proteins and occur on rapid timescales compared to cell growth, MCAS models assume a constant domain size and constant protein amount. This may not always apply. Third, we modeled two-component systems, whereas all known polarity systems have multiple components. More realistic multi-component models of the budding yeast polarity circuit exist (***Goryachev and Pokhilko, 2008***; ***Howell et al., 2009***; ***Kuo et al., 2014***; ***Wu et al., 2015***) and preliminary simulations indicate that they too behave according to the saturation rule. However, adding additional components can yield emergent behaviors not seen in the two-component systems (***Marcon et al., 2016***; ***Otsuji et al., 2010***). Thus, predictions of the saturation rule will need to be tested experimentally to assess whether the insights derived from simple MCAS models are translatable to biological systems.

The most obvious prediction stemming from the saturation rule is that systems should transition between uni- and multi-polarity regimes as total GTPase contents change: lower levels should yield uni-polarity, while higher levels sufficient to allow activator concentrations to approach the saturation point should yield multi-polarity.

In the tractable budding yeast *Saccharomyces cerevisiae*, the master polarity regulatory GTPase, Cdc42, becomes concentrated at polarity sites. Initial peaks of Cdc42 appear to compete on a 1 minute timescale to leave only one winning peak. Moderate overexpression of Cdc42 did not change this behavior (***Howell et al., 2012***). Simultaneous overexpression of Cdc42 and its GEF blocked polarization (***Ziman and Johnson, 1994***), presumably because active GTPase spread throughout the cell cortex. This phenomenon has been explored in Turing models: when component concentrations are too high, the system no longer polarizes, but instead evolves to a stable steady state with high levels of activator uniformly distributed all over the surface (***Howell et al., 2012***).

One way to avoid uniform activation is to increase cell volume as well as total protein content in parallel, maintaining overall concentrations unchanged, which is analogous to the gray line in *Figure 6*. Yeast cells occur naturally as haploids and diploids, and cells with higher ploidy can be constructed. It is also possible to block cytokinesis, generating larger cells due to failed cell division. It appears that cell volume and total protein amount scale with ploidy for most proteins, so that total protein concentrations remain generally unchanged. If we were to keep the activator and substrate concentrations at the homogeneous steady state of an MCAS model constant, then a model with a larger domain size would provide a larger pool of substrate, allowing greater local enrichment of the activator, so that peak activator concentrations would approach the saturation point. This predicts that as cells become larger they should eventually switch from uni- to multi-polarity.

For some filamentous fungi, like *Ashbya gossypii*, development proceeds through a cell enlargement process in which a single shared cytoplasm houses more and more nuclei. This provides a natural system that samples a large range of cell sizes. Cell polarity in *A. gossypii* is thought to be governed by the same Cdc42-centered circuit employed in *S. cerevisiae*, but these cells transition from always having a single polarity site when they are small (following germination), to having two (and then more) polarity sites as they grow larger, leading to hyphal branching (***Knechtle et al., 2003***). Sporadic septation (division separating parts of a single large cell into two smaller ones) can restore a single polarity site to the cell, but continued growth then leads to additional polarity site(s) again. This behavior is consistent with a switch from uni- to multi-polarity according to the saturation rule. A prediction for this system would be that reducing total content of polarity proteins should delay the switch from uni-polar to multi-polar behavior, so that it would take a larger cell to initiate a hyphal branch.

## Conclusions

We have examined the behavior of a family of simplified mathematical models that capture key aspects of the behavior of the Rho-GTPases that regulate the formation of cortical domains in cells. Our analysis suggests that all biologically relevant models of this type (and there are several varieties) display reproducible transitions in system behavior as parameters vary. In particular, each model has a saturation point that depends on model parameters. With low amounts of GTPase, the system forms sharp peaks of active GTPase, but as GTPase levels increase, the peak GTPase concentration approaches the saturation point and the concentration profiles broaden into flat-topped mesas. If there are two or more peaks of GTPase, the peaks will compete with each other until one emerges as the single stable winner. However, the time scale of competition slows dramatically as the peaks broaden, so in practice the systems transition from a situation with rapid cut-throat competition to one in which competition is so languid that peaks co-exist on biologically relevant timescales. Local depletion of the cytoplasmic substrate provides a mechanism of saturation that is universal to all activator-substrate systems, so regardless of the specific biochemical feedback mechanism, a cell that polarizes through local activation and substrate depletion should be able to switch between uni- or multi-polar outcomes by regulating system parameters. The discovery of this intrinsic property of the Rho-GTPase system suggests hypotheses testable in the context of various different cell types.

## Methods

### Model simulation

Simulations of the MCAS models were done on MATLAB with parameters described in the main text. All models were simulated on 1-dimensional domains with fixed spatial resolution of 500 grid points, except the simulations with long *L*, where number of grid points was increased proportionately. The linear diffusion term was implemented by the implicit finite difference method, and the non-linear reaction term by the explicit Euler method every time step. For simulations in the limit *D_v_* → ∞, the mean of *v* was taken every time step. Simulations proceeded with adaptive time stepping according to relative error in the reaction term. The MATLAB code used for simulations is provided in Source Code Files.

### Calculation of competition time

Simulations of competition is generally generated as follows: Two-peak steady states were first generated by simulating the evolution of the homogeneous steady state with an added sine wave. Perturbations were then introduced by increasing the amplitude of the concentration profiles *u*(*x*) *v*(*x*) at regions that we call the first peak by a given percentage (e.g. a 20% increase), and decreasing the amplitude of the second peak by the same percentage (e.g. a 20% decrease). The resulting two unequal peaks were then allowed to compete.

For simulations used in *Figure 4*A and *Figure 5*F, we recorded the measurements of the peak height (*u*_max_) to calculate the saturation index, and the competition time. The steady state *u*_max_ was obtained from the two-peak steady state. The two peaks were then perturbed by increasing the protein content of the left half-domain and decreasing the protein content of the right half-domain, so that each half has 60% and 40% of the original *M*, respectively. For more accurate measurements of the competition time, the two halves were first simulated individually to their own steady states in isolation. Upon the start of competition, the two half-domain were allowed to communicate through diffusion, and the competition time was calculated by measuring the resolution time of two unequal peaks from 60% and 40% at the beginning to 99% and 1%.

### Non-dimensionalization of MCAS Models

We consider the mass-conserved activator-substrate (MCAS) model in a one-dimensional domain with periodic boundary conditions,

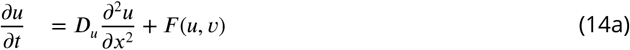

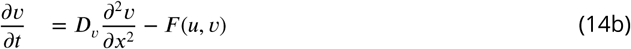

where the diffusion of *v* is much faster than *u*, as set by *D_v_* ≫ *D_u_*.

This model is a mass-conserved version of an activator-substrate model, where *u* is the activator and *v* is the substrate. As the activator concentration *u* increases at certain locations, the substrate *v* is depleted at the same rate, therefore, the mass *M* is a constant conserved for all time,

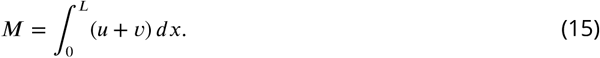

The reaction terms of these models, given by *F*(*u, v*), generally contain a *v*-dependent activation term with nonlinear positive-feedback, and a *v*-independent inactivation term.

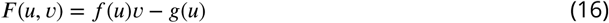

For example, the model proposed by ***Mori et al.*** (***2008***). has an *f*(*u*) with a saturable non-linear term and a linear *g*(*u*),

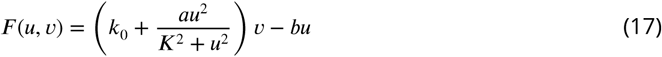

We assume negligible *k*_0_, and rewrite this system with dimensionless variables 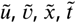 by scaling the length by the domain size *L*, and time by *T* and *u* and *v* by *U*,

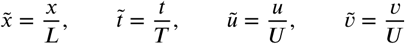

yielding

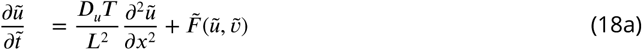

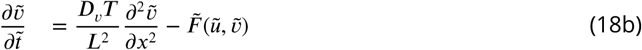

where the nondimensionalized reaction terms are now

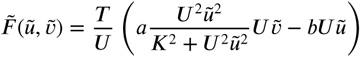

and the non-dimensional mass 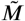 is

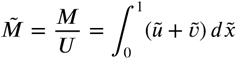

Setting the timescale and concentration scale as

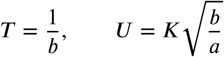

and dropping the tildes, puts the system into the form

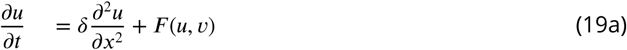

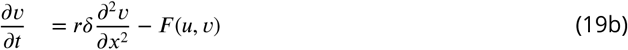

where

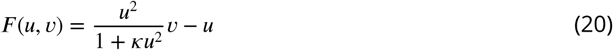

and the dimensionless parameters are

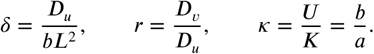

Setting κ to zero, we obtain a “Turing-type” system with

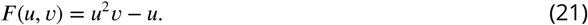

### Steady state solutions for the *D_v_* ≫ *D_u_* limit

We first consider the simplified case where *r* → ∞. The solutions *u, v* are expanded as regular perturbation series with respect to inverse powers of *r*,

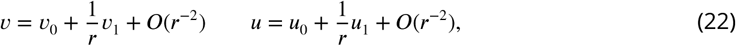

and substituted into (19ab). The leading order equation for *u*_0_ at *O*(1) mirrors (19a),

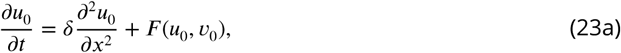

while at *O*(*r*), the leading order equation for *v*_0_ becomes *v*_0*,xx*_ = 0. Subject to the periodic boundary conditions, this forces *v*_0_ to be spatially uniform, but it can depend on time, *v*_0_ = *v*_0_(*t*). In order to obtain an equation defining the evolution of *v*_0_ we proceed to the next term in the expansion. At *O*(1) we find that in order for a solution for *v*_1_ to exist, the inhomogeneous terms in the equation must satisfy a solvability condition. Set by the Fredholm alternative theorem, this condition gives the evolution for *v*_0_(*t*) in terms of the reaction rate averaged over the domain,

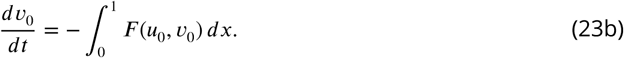

This equation effectively describes the evolution of the average substrate concentration in the well-mixed limit. Moving forward, we will drop the zero-subscripts and focus on solving this leading order system.

At steady state, the solution (*u_ss_*(*x*)*, v_ss_*) satisfies the system of equations

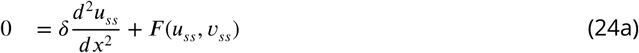

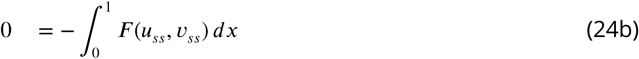

and the constraint on the total mass

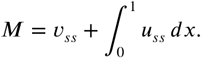

To understand the properties of the solutions it is helpful to integrate (24a) with respect to *du* to obtain

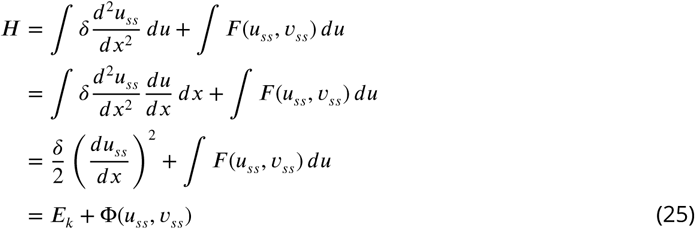

where *H* is a constant and

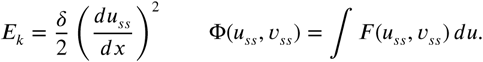

Solutions *u_ss_*(*x*) can be found that have any number of peaks, *n*, with corresponding spatial period 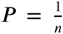. These steady states are spatially periodic multi-peak solutions, with all peaks being all *n* identical and are equally spaced within each solution (*Figure 9*-Figure supplement 1).

**Figure 9.**
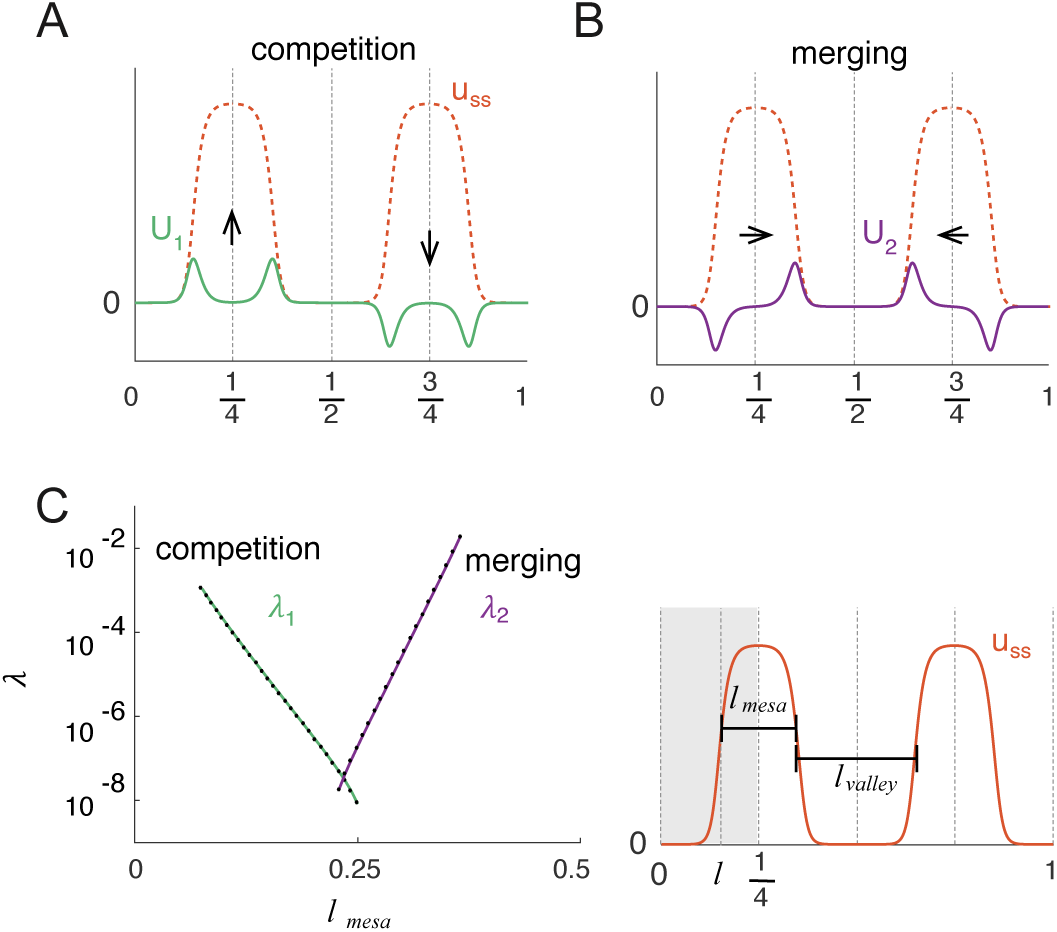
Linear Stability Analysis reveals two unstable modes: competition and merging. The two eigenmodes *U*_1_ and *U*_2_ constructed from the linear stability analysis represents competition (A) and merging (B) of the two peaks in the full domain. C) The eigenvalue for competition *λ*_1_ decreases exponentially with increasing peak width *ℓ*_mesa_, and the eigenvalue for merging *λ*_2_ increases exponentially with decreasing distance between peaks *ℓ*_valley_. The eigenvalues for competition and merging were calculated by the shooting method for each steady state solution with varying *M* from 10 to 32 at an increment of 0.5. The lengths of the valley and the mesa are equivalent to 2*ℓ* and 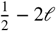, with *ℓ* defined as the position of half max. **Figure 9-Figure supplement 1.** An MCAS system with a given set of parameters can yield solutions of different periodicities *n*. **Figure 9-Figure supplement 2.** Construction of the two forms of the eigenmode *U* (*x*) using even and odd extensions of the quarter-domain solutions *U*_1_(*x*) (A) and *U*_2_(*x*) (B) for a two-peak solution *u_ss_*(*x*). **Figure 9-Figure supplement 3.** Schematic representations of the shooting method constructions for the *λ*_1_ (shooting from 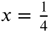) (A), and the *λ*_2_ (shooting from 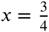 to 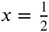) eigenmodes (B), which represent competition and merging modes respectively. **Figure 9-Figure supplement 4.** Construction of the approximate *U*_1_(*x*) eigenfunction using hyperbolic functions with a midpoint boundary condition (36) at position *ℓ*.

The local extrema, *u*_min_ and *u*_max_, occur where *du_ss_*/*dx* = 0. A direct consequence of this is that for a given value of *H*, the integral of *F*(*u, v*) from *u*_min_ to *u*_max_ must be zero to satisfy (25). This condition has been referred to as the **wave-pinning condition** (Mori et al., 2008), and is general to all MCAS models,

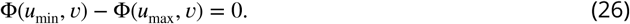

There also exists a special value of the cytoplasmic concentration *v*, which we call *v*_sat_ at which

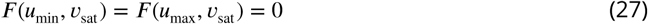

We refer to this condition as the **saturation condition**, which is crucial for later discussion of competition time scale. At saturation, *u*_min_ = 0 and we label the value of *u*_max_ as *u*_sat_, which is the largest value of *u*_max_ possible.

### Stability Analysis of Multipeak Steady States

The key question of whether competition happens between two peaks can be answered mathematically by assessing the stability of the two-peak steady state solution (*u_ss_*(*x*)*,v_ss_*). Consider the multi-peak steady state when peak number *n* = 2, a steady state solution (*u_ss_*(*x*)*,v_ss_*) has two identical peaks centered at 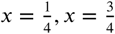. Each peak is reflectionally symmetric about its maximum and the overall solution is also symmetric about 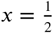 (see *Figure 9*-Figure supplement 2.).

The stability of the two-peak solution is studied by assuming small perturbations of the form

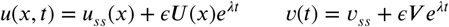

where *λ* and (*U* (*x*)*, V*) satisfy the linearized eigenvalue problem

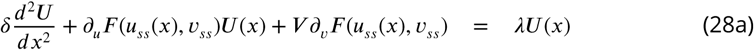

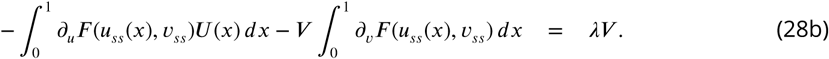

The solution (*u_ss_*(*x*)*, v_ss_*) is unstable if there exists at least one eigenvalue with Re(*λ*) *>* 0.

To approach this question it is sufficient to restrict attention to a particular form of eigenmode, one with *U*(*x*) being antisymmetric with respect to 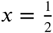, namely *U*(*x* + 1/2) = −*U*(*x*). Since *u_ss_*(*x*) is symmetric with respect to 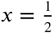, the integrand of the first integral in (28b) is antisymmetric and hence the integral on the whole domain must vanish. Consequently, for this type of eigenmode, *V* = 0 solves (28b) and the system reduces to

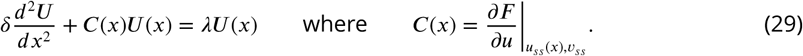

To solve (29) on the full domain *x* ∈ [0, 1] with periodic boundary conditions, it is sufficient to solve the equation on a quarter domain, 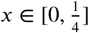, with boundary conditions

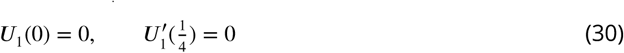

or

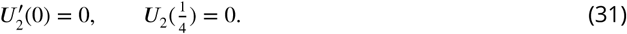

The periodic function *U*(*x*) is then constructed by extending *U*_1_(*x*) symmetrically with respect to 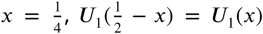 (*Figure 9*-Figure supplement 3A), or by similarly extending *U*_2_(*x*) anti-symmetrically, 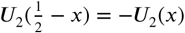 (*Figure 9*-Figure supplement 3B).

We give a shooting argument to show that there is a positive eigenvalue *λ*_1_ *>* 0 for solutions having the form given by (30). Differentiating the steady state equation (24a) with respect to *x*, we obtain

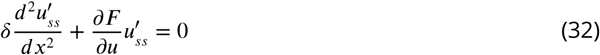

Therefore, 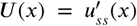 solves (29) with *λ* = 0. Note that in this case 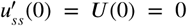 and 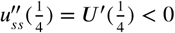 (there is a finite curvature at *u*_max_) and hence the second condition in (30) is not satisfied. If *λ >* max(*C*(*x*)), we can re-write (29) as *δU"* = (*λ*−*C*(*x*))*,U >* 0 and *U*(*x*) will be monotonely growing and 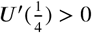. Since we have constructed solutions achieving positive and negative values 4 for 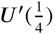, by continuity, there exists an eigenvalue in the range 0 < *λ*_1_ *<*max(*C*(*x*)) that will yield an eigenmode satisfying (30) (*Figure 9*-Figure supplement 4B). Similarly, there exists a second eigenmode of the *U*_2_(*x*) (31) form with eigenvalue in the range 0 *< λ*_2_ < max(*C*(*x*)) *Figure 9*-Figure supplement 4B).

To summarize, there exist *λ*_1_*, λ*_2_ *>* 0, thus the two-peak steady state is not stable. Further, the eigenfunction *U*_1_ corresponds to one peak growing and the other shrinking, i.e. competition (*Figure 9*A). The eigenvalue *λ*_1_ corresponds to the timescale for competition. On the other hand, the eigenfunction *U*_2_ corresponds to neighboring sides of each peak growing while the other sides shrink such that peaks merge with each other. The eigenvalue *λ*_2_ corresponds to the timescale for merging (*Figure 9*B).

### The eigenvalues for competition and merging

As two-peak steady states are always unstable, the distinction between competition and co-existence of two peaks (*Figure 3*C-E) does not reflect a change in stability, but rather, a change in the time scale on which competition occurs. As it has been reported that mesas are meta-stable, we inquired how the eigenvalues of competition and merging change with increasing width of the peaks in a two-peak steady state. We will show that

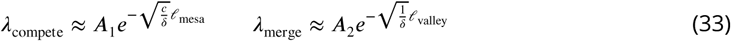

where δ is the diffusion constant of *u*, and *A*_1_*, A*_2_*, c* are constants. *ℓ*_mesa_ and *ℓ*_valley_ are defined as in *Figure 9*C with *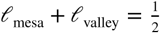*. Our derivation proceeds in the following steps:

1. Approximate the steady state as a step function for *C*(*x*) in (29), one segment of which is approximated using *u*_min_ for and the other using *u*_max_. The two segments are connected with a mid-point boundary condition at *x* = *ℓ* (*Figure 9*-Figure supplement 4).
2. Calibrate the mid-point boundary condition using the translational eigenmode 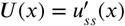. *λ* = 0.
3. Solve the approximated version of (29) for the competition eigenvalue subject to the competition boundary condition (30) and the mid-point boundary condition calculated in the previous step.
4. Similarly, solve (29) for the merging eigenvalue subject to the merging boundary condition (31).

#### Approximating the steady state solution

Without loss of generality, let *F*(*u, v*) be the non-dimensionalized wave-pinning model (20),

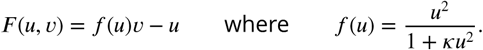

For near-saturation two-peak solutions that have a mesa shape, *v* is chosen such that (27) is met, thus at *u* = *u*_max_,

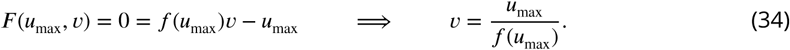

We try to approximate the eigenvalue problem (29) on a quarter domain by a step function. The two segments *U*(*x*) and 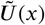 satisfy (29) with *u_ss_*(*x*) approximated by *u*(*x*) = *u*_min_ and *u*(*x*) = *u*_max_ respectively. The length of *U* and 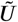 are the length of the valley 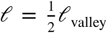 and the width of the mesa 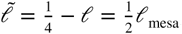, respectively. As *f′*(*u*_min_) = 0, we can re-write (29) into the step function

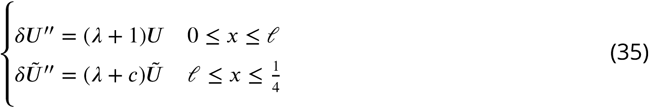

where

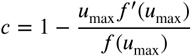

with the **mid-point boundary conditions**

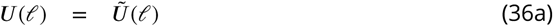

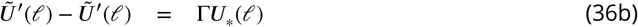

#### Solving the mid-point boundary condition

Since 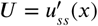 is the translationa leigen mode for *λ*= 0, we first use this solution and (35) to determine Γ in (36b). The piecewise solutions of (35) are given by

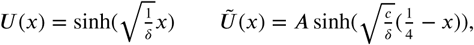

and then (36) takes the form

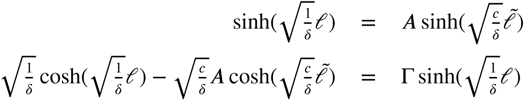

yielding

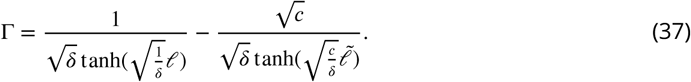

#### Approximating the competition eigenvalue

To satisfy the **competition boundary condition** (30), the solutions *U,* 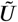 are chosen in (35) as

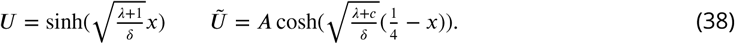

Then midpoint boundary condition (36b) can be rewritten with Γ as

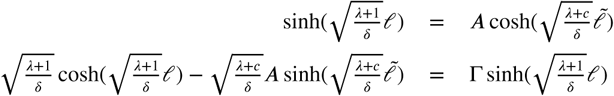

This yields the equation

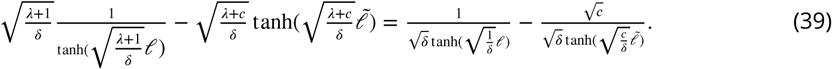

After rearranging terms and making use of simplifications for small *δ*, this equation can be reduced to

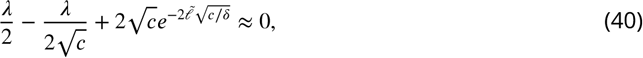

which finally yields

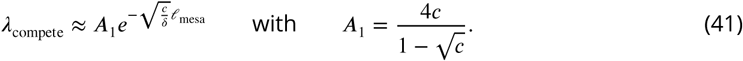

#### Approximating the merging eigenvalue

Similarly, to satisfy the **merging boundary condition** (31), the solutions *U,* 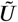are chosen as

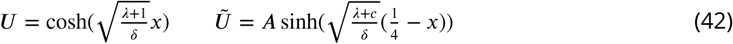

Substituting these solutions into the midpoint boundary conditions (36b) yields

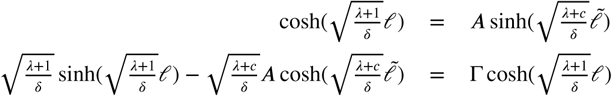

This system of equations can then be reduced to the condition

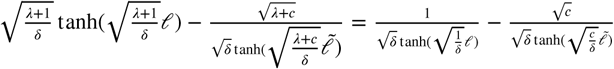

Again, neglecting smaller terms in the limit that *δ* is small, we obtain

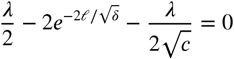

which finally yields

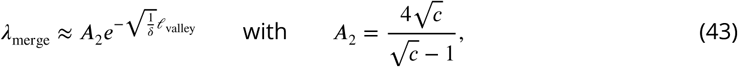

and recall that 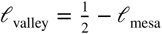. We can numerically calculate two-peak steady-states over a range of values for *ℓ*_mesa_ by varying the total mass *M*. The computed results for *λ*_compete_ and *λ*_merge_ for the two-peak steady states confirm these analytical predictions (*Figure 9*C).

The width of a mesa is an indicator that perfectly correlates with how close to saturation a peak is. Dedining a saturation index as (*u*_sat_ − *u*_max_)/*u*_sat_, and *ℓ*_mesa_ as normalized by 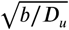, we find that two peak steady states of the dimensional model (***Equation 5***) plotted in *Figure 4* collapse into a perfect correlation, no matter what parameter we change (*Figure 4*-Figure supplement 1). The relationship between *ℓ*_mesa_ shows that the wider the mesa, the more saturated the two peak steady states are, and thus the less efficient competition will be.

### Turing stability of the Wave-pinning model

We investigate the stability of the wave-pinning model below, and find that with appropriate parameters, the Wave-pinning model is indeed Turing unstable.

The reaction term of the wave-pinning model (20) has three roots. One is the trivial solution, which is always Turing stable:

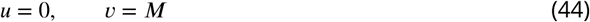

The non-trivial solutions can be obtained from

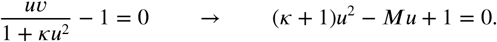

This yields two solutions

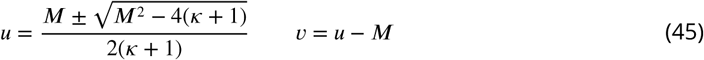

under the condition

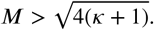

The condition for Turing instability in MCAS models reads as follows (***Mori et al., 2008***; ***Rubinstein et al., 2012***):

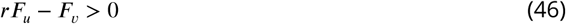

where *r* = *D_v_*/*D_u_* and *F_u_* and *F_v_* are the derivatives of *F*(*u, v*) with respect to *u*,

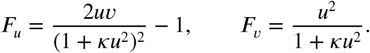

Since the homogeneous steady states satisfy

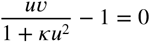

then condition (46) becomes

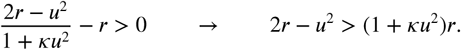

Therefore, the Turing unstable condition in the non-dimensional system (19, 20) reads:

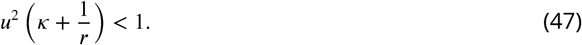

In the Turing-type model at when *k* is small, the system is easily Turing unstable due to large ratio *r* between the two diffusion constants.

## Author Contributions

J.-G.C. and D.J.L. developed the concept of this article; J.-G.C. developed the model and conducted the simulations; J.-G.C., T.C.E., T.P.W., and D.G.S. developed the formal mathematical analysis; J.-G.C. and D.J.L. drafted the manuscript; All authors contributed in reviewing and editing the manuscript.

## Acknowledgments

We thank Amy Gladfelter, Stefano Di Talia, Sam Ramirez, Ben Woods, and all members of the Lew lab for stimulating discussion and comments on the manuscript. This work was supported by NIH/NIGMS grant GM62300 to D.J.L.

---

**Figure 4-Figure supplement 1.**
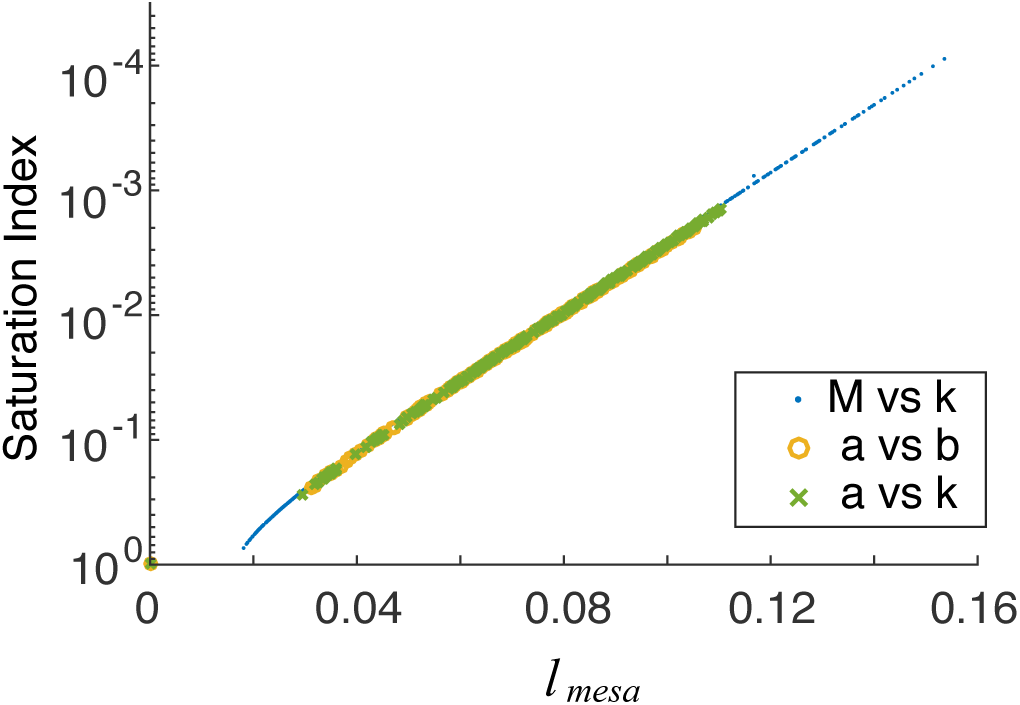
Peak width, *ℓ*_mesa_, is a robust indicator of saturation over a broad range of system parameters. Data points collected from simulations in *Figure 4*A.

**Figure 9-Figure supplement 1.**
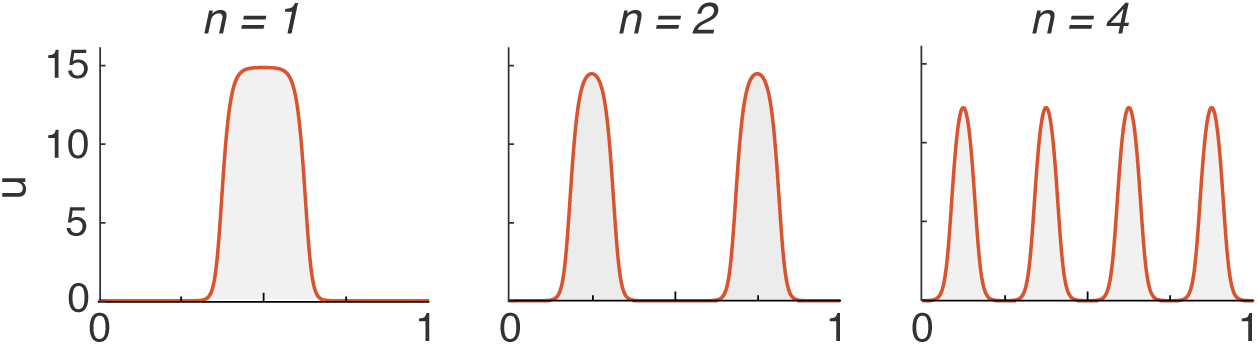
An MCAS system with a given set of parameters can yield solutions of different periodicities *n*.

**Figure 9-Figure supplement 2.**
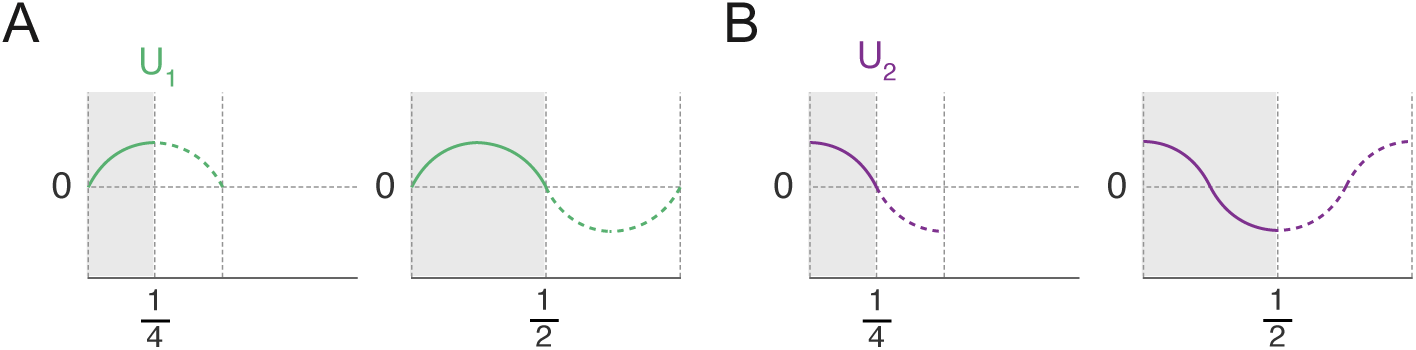
Construction of the two forms of the eigenmode *U* (*x*) using even and odd extensions of the quarter-domain solutions *U*_1_(*x*) (A) and *U*_2_(*x*) (B) for a two-peak solution *u_ss_*(*x*).

**Figure 9-Figure supplement 3.**
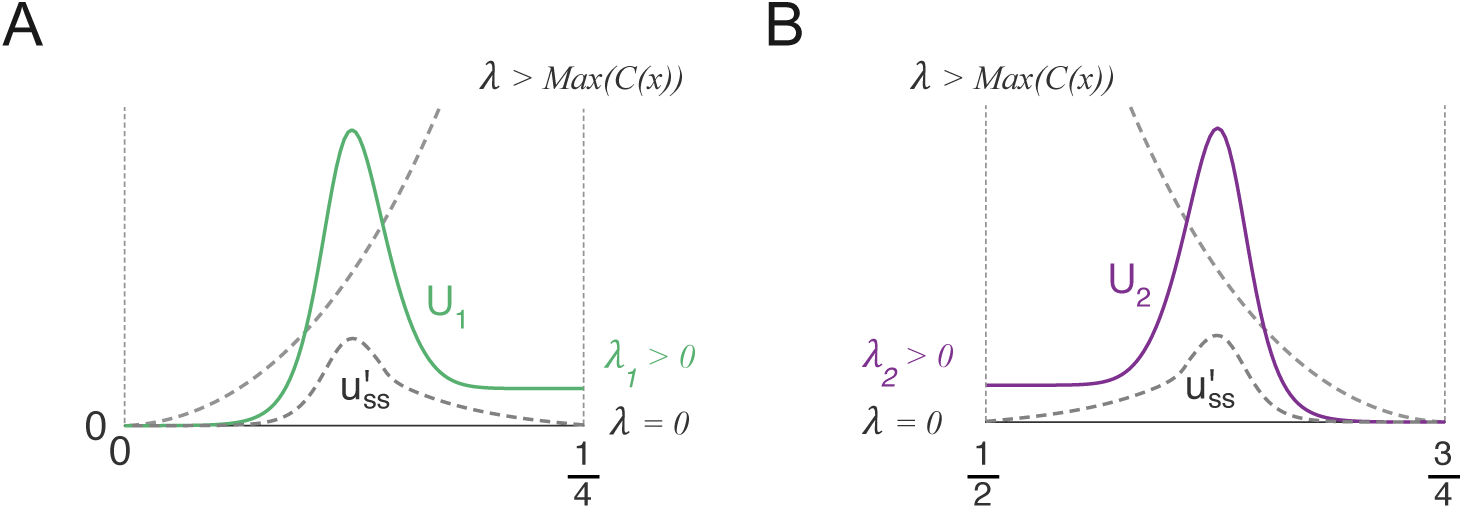
Schematic representations of the shooting method constructions for the *λ*_1_ (shooting from 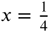) (A), and the *λ*_2_ (shooting from 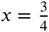 to 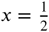) eigenmodes (B), which represent competition and merging modes respectively.

**Figure 9-Figure supplement 4.**
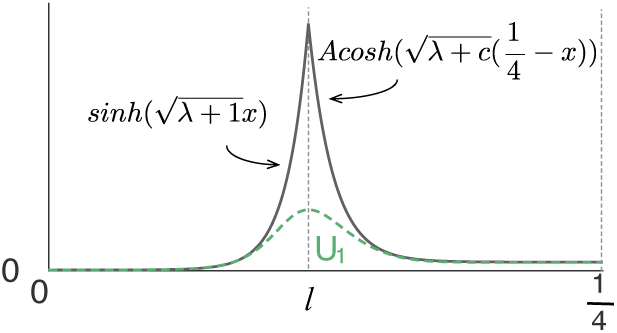
Construction of the approximate *U*_1_(*x*) eigenfunction using hyperbolic functions with a midpoint boundary condition (36) at position *ℓ*.

